# Structural basis of human 20S proteasome biogenesis

**DOI:** 10.1101/2024.08.08.607236

**Authors:** Hanxiao Zhang, Chenyu Zhou, Zarith Mohammad, Jianhua Zhao

## Abstract

New proteasomes are produced to accommodate increases in cellular catabolic demand and prevent the accumulation of cytotoxic proteins. Formation of the proteasomal 20S core complex relies on the function of the five chaperones PAC1-4 and POMP. To understand how these chaperones facilitate proteasome assembly, we tagged the endogenous chaperones using CRISPR/Cas gene editing and examined the chaperone-bound complexes by cryo-EM. We observed an early α-ring intermediate subcomplex that is stabilized by PAC1-4, which transitions to β-ring assembly upon dissociation of PAC3/PAC4 and rearrangement of the PAC1 N-terminal tail. Completion of the β-ring and dimerization of half-proteasomes repositions critical lysine K33 to trigger cleavage of the β pro-peptides, leading to the concerted dissociation of POMP and PAC1/PAC2 to yield mature 20S proteasomes. This study reveals structural insights into critical points along the assembly pathway of the human proteasome and provides a molecular blueprint for 20S biogenesis.

## BACKGROUND

The proteasome system is responsible for degrading the majority of proteins in eukaryotic cells and plays a key role in metabolism and protein quality control mechanisms (1–8). Proteasome function is crucial for maintaining nutrient balance and removing cytotoxic polypeptides that can cause cellular stress and dysfunction. Consequently, decreases in proteasome levels and activity are associated with aging and neurodegenerative diseases such as Alzheimer’s disease and Parkinson’s disease (9–12). To maintain homeostasis, cells must replenish existing proteasomes as well as produce new proteasomes in response to elevated levels of damaged and misfolded proteins (5, 6, 13, 14).

The production of new proteasomes is an intricate process that requires the precise assembly of many different components and involves multiple chaperones. Early in the biogenesis pathway, proteasome assembly chaperones (PAC) 1-4 coordinate the arrangement of 7 different α subunits (α1-7) into a hetero-heptameric α-ring (15–17). Next, the proteasome maturation protein (POMP) chaperone assembles the 7 different β subunits onto the α-ring (18–21) with concomitant unbinding of the PAC3/PAC4 heterodimer. The completed α-ring/β-ring heterocomplex (half-proteasome) dimerizes to form the pre-20S complex, which activates the autocatalytic cleavage of the β pro-peptides (22, 23). Finally, POMP and the PAC1/PAC2 heterodimer dissociate to yield the mature 20S proteasome (15, 18, 24). Recent structural work has shed light on the mechanisms underlying β-ring assembly by POMP and PAC1/PAC2 during the middle stages of 20S biogenesis in yeast (25). On the other hand, the mechanisms underlying the early and late stages of the assembly process remains poorly understood.

The complexity of proteasome assembly has made it challenging to replicate the process in vitro. Knock-down and mutation of some subunits in yeast and mammalian cells have been useful strategies in trapping certain proteasomal intermediates for biochemical and structural analyses (18, 24–26). However, decreases in cellular fitness and changes in complex stoichiometry associated with traditional approaches have complicated the in-depth structural characterization of the proteasome biogenesis pathway (15, 16, 27–30). We have previously developed efficient strategies of tagging and purifying endogenous proteins for cryo-EM analysis (31, 32), which we have employed in this study to dissect the stepwise assembly pathway of the human 20S proteasome. Here, we provide a broad structural analysis of 20S biogenesis, which clarifies the role of each chaperone across the assembly process and reveals new molecular insights into PAC3/PAC4 function and late-stage maturation of the pre-20S complex.

## RESULTS

### Overall assembly of the 20S proteasome

Assembly of the proteasomal 20S core complex is orchestrated by five dedicated chaperones. To study their function, we individually tagged three of the five 20S chaperones in Expi293F cells using CRISPR/Cas gene editing (31, 33) and separately purified the endogenous chaperone-associated complexes. Analysis of the samples by single particle cryogenic electron microscopy (cryo-EM) enabled structure determination of intermediate complexes at each major stage of 20S assembly (Figure 1A, Supplementary Figures 1-3, Supplementary Table 1). Overall, each α and β subunit interacts with at least one of the 20S chaperones except for β7 (Figure 1B). Evidence of protein dynamics can be observed throughout the assembly process. In particular, the α subunits undergo substantial movements that appear to correlate with binding of the β subunits (Figure 1C). On the other hand, the chaperones mainly undergo local structural rearrangements with limited changes in overall architecture (Figure 1D). In the following, we describe mechanistic insights into each of the major stages of 20S assembly.

**Figure 1.**
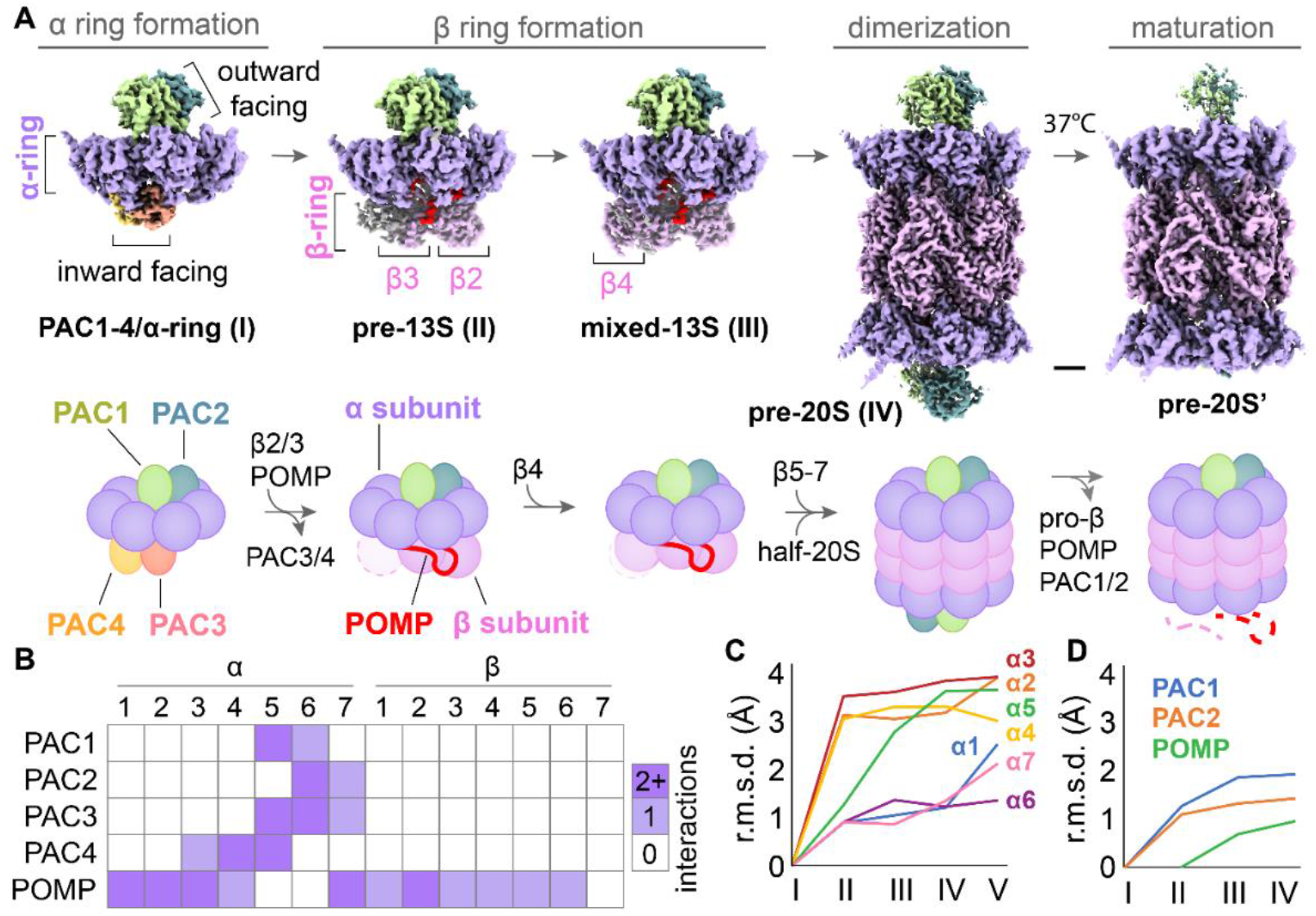
Stepwise assembly of the 20S proteasome is mediated by multiple chaperone interactions. (**A**) Cryo-EM structures of human 20S intermediate complexes depicting the stepwise assembly and maturation of the proteasome. Assembly begins with formation of the α-ring, aided by PAC1-4. The β-ring is formed on the α-ring, aided by POMP. Dimerization of α-ring/β-ring half-proteasomes triggers cleavage of the β pro-peptides (pro-β) and dissociation of POMP and PAC1/PAC2 to yield the mature 20S proteasome. Scale bar: 20 Å. (**B**) Interaction map between proteasome chaperones and 20S subunits. Each chaperone makes extensive contacts with multiple 20S subunits, with PAC1-4 interacting primarily with the α4-7 subunits while POMP interactions are spread out across many α and β subunits. (**C**) Root mean square deviation (r.m.s.d.) values of the α subunit atomic positions across the PAC1-4/α-ring (I), pre-13S (II), mixed-13S (III), pre-20S (IV), and mature 20S (V, PDB 7NAN) structures. (**D**) R.m.s.d. values of proteasome chaperone atomic positions across the PAC1-4/α-ring (I), pre-13S (II), mixed-13S (III), and pre-20S (IV) structures.

### α-ring formation

20S proteasome assembly begins with formation of the α-ring (*34*) and involves PAC1-4. To gain molecular insights into α-ring formation, we tagged PAC3 (PSMG3) in Expi293F cells and purified the PAC3-associated complexes. Cryo-EM analysis of the sample resulted in a structure of the α-ring complex bound by PAC1-4 at ~3.0 Å resolution (Figure 2A, Supplementary Figures 2A and 3). While the PAC1/PAC2 and PAC3/PAC4 heterodimers bind on different sides of the α-ring, these two heterodimers interact with overlapping sets of α subunits. α5 and α6 make the most extensive contacts with PAC1-4, with the N termini of α5 and α6 inserting into a hydrophobic crevice formed between PAC1 and PAC2 (Supplementary Figure 4A), which was also observed in yeast (*25, 35*). Meanwhile, PAC3 and PAC4 clamp onto one of the α-helices in α5, explaining the tight binding of PAC3 and PAC4 to α5 (Supplementary Figure 4B) (*28, 36, 37*). PAC3 also interacts with α6 (Supplementary Figure 4C). Additionally, the C terminus of PAC1 inserts into the HbYX-binding pocket formed between α5 and α6 (Supplementary Figure 4D) (*38, 39*). The extensive structural contacts between PAC1-4 and α5-6 suggest that these components may form an initial subcomplex that catalyzes α-ring formation.

**Figure 2.**
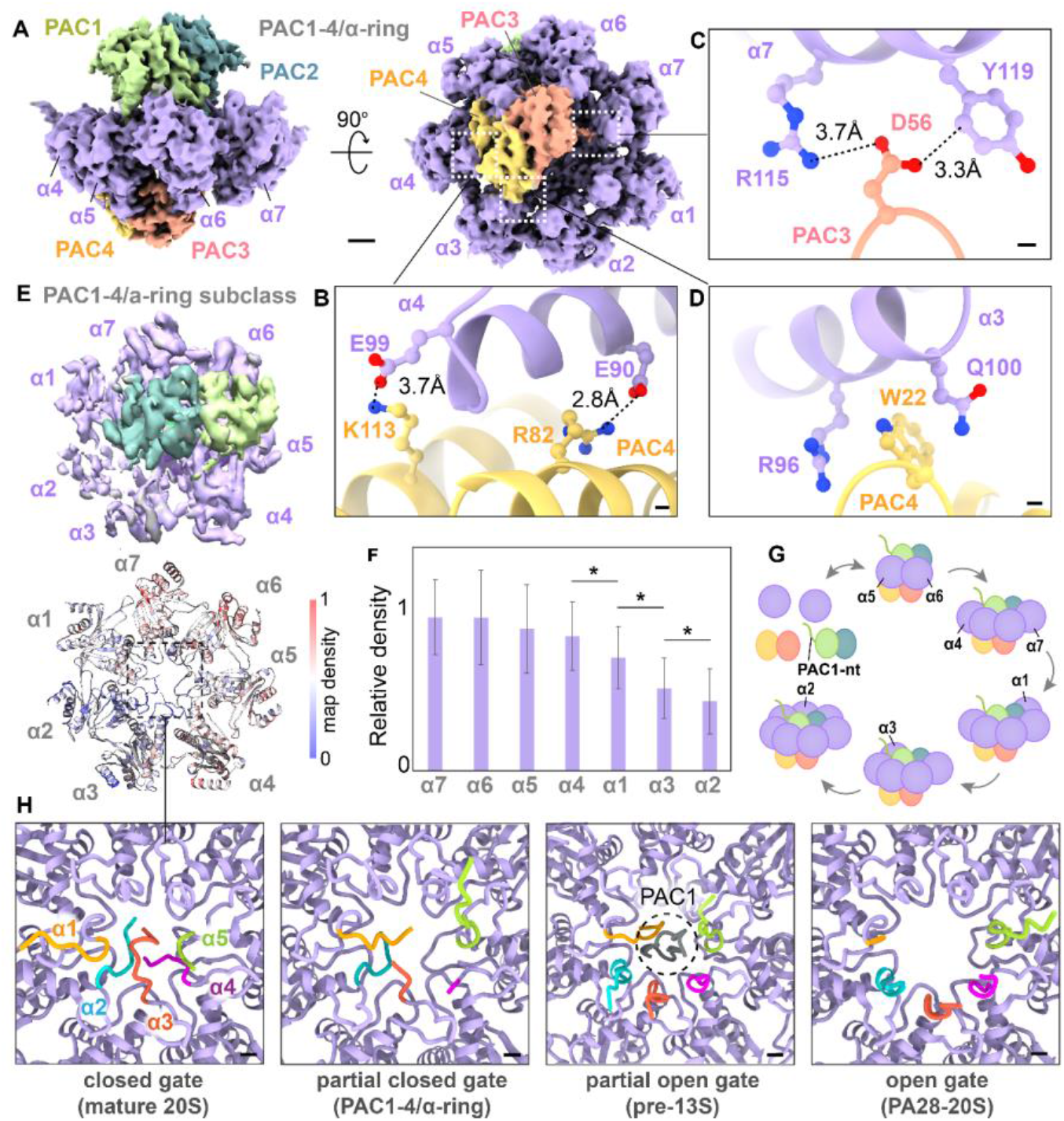
PAC1-4 helps to stabilize the α4-7 subcomplex to initiate proteasome assembly. (**A**) Cryo-EM structure of the PAC1-4/α-ring complex showing the distinct binding sites of PAC1/PAC2 and PAC3/PAC4. PAC1/PAC2 bind to α5-7 on the outward facing side of the α-ring while PAC3/PAC4 bind to α4-6 on the concave inward facing side of the α-ring. Scale bar: 15 Å. (**B**) PAC4 K113 and R82 form salt bridges with α4 E99 and E90, respectively. Scale bar: 5 Å. (**C**) PAC3 D56 forms a salt bridge with α7 R115 and coordinates with α7 Y119. Scale bar: 5 Å. (**D**) PAC4 W22 fits into a pocket formed by α3 R96 and Q100. Scale bar: 5 Å. (**E**) PAC1-4/α-ring subclass from 3D classification of cryo-EM data showing sub-stoichiometric binding of α-subunits. (**F**) Quantification of the density values for each α-subunit reveals lower map densities for α1-3. *p<0.001. (**G**) Schematic depicting α-ring assembly. (H) Positions of the α subunit N-terminal tails in a closed gate (mature 20S, PDB 7NAN), partially closed gate (PAC1-4/α-ring), partially open gate (pre-13S), and fully open gate (PA28-20S, PDB 7NAO). The N-terminal tail of PAC1 is observed in the pre-13S structure but not in the PAC1-4/α-ring structure.

To gain further insights into α-ring formation, we carried out computational 3D classification analysis of the PAC1-4/α-ring cryo-EM data. The analysis revealed a 3D subclass with strong map densities for α4-7 and lower map densities for α1-3 (Figure 2E, Supplementary Figure 2A). In contrast, similar analysis of β-ring assembly intermediates revealed consistent α subunit map densities (Figure 5A). This result indicates that PAC1-4/α4-7 likely forms a stable subcomplex (*40*). PAC1-4/α4-7 is stabilized by multiple contacts between α4 and PAC4, including salt bridges between α4 E90 and PAC4 R82, and between α4 E99 and PAC4 K113 (Figure 2B). Meanwhile, α7 makes contacts with both PAC2 and PAC3, with the C-terminal tail of PAC2 inserting into the HbYX-binding pocket formed between α6 and α7 (Supplementary Figure 4D). Additionally, a loop in PAC3 extends out toward α7 to form a salt bridge between α7 R115 and PAC3 D56 (Figure 2C). While PAC1-4 are crucial for stabilizing α4-7, they do not appear to play a substantial role in the incorporation of α1-3 with the only contact being PAC4 W22 stacking in between α3 R96 and Q100 (Figure 2D). α4-7 have the highest map densities in the 3D subclass, followed by α1, α3, and α2 (Figure 2F, Supplementary Figure 4E). This result suggests that α1 is likely added first after formation of the PAC1-4/α4-7 subcomplex, followed by incorporation of α3, and then α2 is inserted last to complete the α-ring (Figure 2G). These structural findings are consistent with results from knockdown experiments showing α2 incorporates after α1 (*40*).

Substrate entry into the proteasome is regulated by the N-terminal tails of the α subunits, which form a closed gate in the mature 20S complex (Figure 2H). In structures of β-ring assembly intermediates, the N-terminal tail of PAC1 is inserted into the α-ring pore and blocks the open gate (Figure 2H, Supplementary Figure 5C). It has been postulated that the N-terminal tail of PAC1 may help to coordinate α-ring assembly in yeast (*25, 41*). Surprisingly, the N-terminal tail of PAC1 is absent from the α-ring pore in the human PAC1-4/α-ring structure reported here (Figure 2H, Supplementary Figure 5C). The N-terminal tails of some α-subunits in the PAC1-4/α-ring assembly adopt unique conformations that result in a partially closed gate (Figure 2H). Specifically, the N-terminal tails of α2-4 are oriented inwards in the PAC1-4/α-ring complex, similar to the mature 20S closed gate conformation. The unique conformations of the α2 and α3 N-terminal tails in the PAC1-4/α-ring complex would sterically clash with the N-terminal tail of PAC1 if it was inserted in the pore (Supplementary Figure 5C). Consequently, the N-terminal tail of PAC1 is inserted only after α-ring formation is completed.

### β-ring formation

The assembly of the β-ring is initiated by the binding of the POMP chaperone and β2 to the completed α-ring. Subsequently, PAC3/PAC4 must dissociate from the α-ring to make room for the incoming β subunits. To gain insight into how this occurs, we tagged POMP in Expi293F cells and purified the POMP-associated complexes. Cryo-EM analysis of the sample revealed a mixture of β-ring assembly intermediates (Supplementary Figure 2B), which were separated into two major classes using computational 3D classification of the cryo-EM images. These two structures correspond to a pre-13S complex with partial binding of β4 at ~3.0 Å resolution and a mixed-13S complex with partial binding of β1, β5, and β6 at ~3.0 Å resolution (Supplementary Figure 3) (*42*). Interestingly, comparison of the PAC1-4/α-ring and pre-13S structures shows that POMP would fit into the concave surface on the α-ring next to PAC3/PAC4 with minimal overlap between the structures (Figure 3A). Additionally, β2 binds to the α-ring on the opposite side of where PAC3/PAC4 is bound. Steric clash would occur in one small one region of PAC4 with the C-terminal tail of POMP and the β2 pro-peptide (Figure 3B-C). The structural arrangement of the POMP C terminus and β2 pro-peptide appears to be stabilized by interactions with β3, which include a salt bridge between β3 R66 with POMP E125. The release of PAC3 is coupled to the binding of β3 (*24*), and our structural data provides a possible mechanism of how this may occur. POMP and β2 binds to the PAC1-4/α-ring complex and recruit β3, which induces structural changes in the POMP C terminus and β2 pro-peptide (Figure 3E). These changes displace PAC4, which would cause the concurrent dissociation of PAC3.

**Figure 3.**
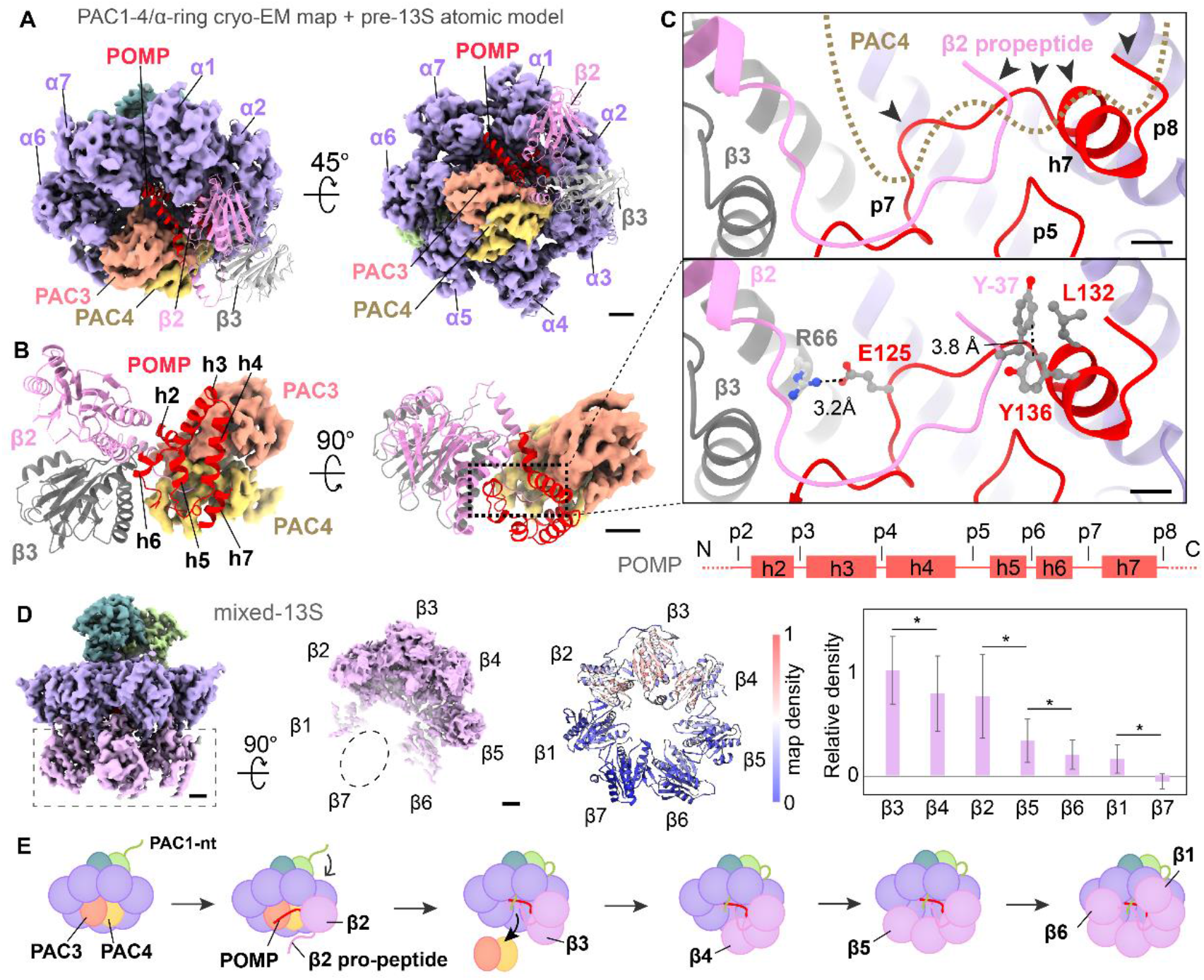
POMP and β2 displaces PAC3/PAC4 to enable β-ring assembly. (**A**) Cryo-EM map of PAC1-4/α-ring overlayed with the pre-13S atomic model. POMP fits into the space between PAC3/PAC4 and α1-2. β2 and β3 bind to the α-ring on the opposite side of PAC3/PAC4. Scale bar: 15 Å. (**B**) Isolated views of the overlayed structures show good fit between POMP and PAC3/PAC4 with little overlap between the chaperones. Scale bar: 15 Å. (**C**) The PAC4 structure clashes with the C-terminal region of POMP and the N-terminal tail of the β2 pro-peptide (arrowheads), which are stabilized by interactions with β3. Scale bar: 10 Å. (**D**) Cryo-EM map of the mixed-13S structure shows sub-stoichiometric binding of the β subunits. Quantification of the density values for each subunit reveals lower map densities for β5, β6, and β1. No map density above background is observed for β7. *p<0.001. Scale bar: 18 Å. (**E**) Schematic depicting β-ring assembly.

The pre-13S and mixed-13S cryo-EM maps represent averages of multiple β-ring assembly intermediates (Figure 3D, Supplementary Figure 2B). Therefore, we may infer the order of β subunit incorporation from their cryo-EM map density levels. The pre-13S cryo-EM map shows strong density for β2/β3 and lower density for β4 (Figure 2B, Supplementary Figure 5B), indicating that β4 is added after β3 as expected (*24*). In the mixed-13S map, β2-4 have comparable densities that are higher than for the other β subunits (Figure 3D, Supplementary Figure 6). β3 has slightly higher average map density compared to β2 and β4, which reflects the flexibility of the terminal β subunits of the incomplete β-ring (Figure 3D, Supplementary Figures 3E, 6). After β4, β5 is likely incorporated next, followed by β1 and β6 (Figure 3D-E). Interestingly, no density above background is observed for β7 in either the pre-13S or mixed-13S cryo-EM maps. This indicates that β7 is the last subunit added to yield the half-proteasome (*24, 43, 44*) and two half-proteasomes will quickly dimerize to form the pre-20S complex.

### Dimerization and autocatalytic activation

The catalytic β subunits (β1, β2, β5) are expressed with N-terminal pro-peptides (Supplementary Figure 1B-C) that are cleaved after formation of the pre-20S complex. To gain insight into how dimerization activates the catalytic β subunits, we tagged PAC2 (PSMG2) in Expi293F cells and purified the PAC2-associated complexes. Cryo-EM analysis of the sample resulted in a structure of the pre-20S complex at ~2.8 Å resolution (Figure 4, Supplementary Figures 2C, 3). The cryo-EM map shows clear density for PAC1/PAC2 and POMP, as well as an unknown strand of density located at the interface between the two β4 subunits (Supplementary Figure 7). The close proximity of this density to threonine T1 of β5 suggests it may be part of the β5 pro-peptide (Supplementary Figure 1C), which is crucial for half-proteasome dimerization and can work in trans (*24, 45, 46*). However, we cannot rule out the possibility that the density may also represent the N terminus of POMP (*19*). Comparison of the β pro-peptide densities reveals that the different pro-peptides are processed at different rates. The β1 pro-peptide density appears to be lower compared to the β2 pro-peptide (Figure 4E, Supplementary Figure 8), although the difference is not statistically significant. Interestingly, little density for the β5 pro-peptide can be observed near β5 T1, indicating that the β5 pro-peptide is mostly cleaved and has dissociated in the pre-20S cryo-EM map reported here (Supplementary Figure 8). These data indicate that the β5 pro-peptide is likely processed first after dimerization (Figure 4F), followed by processing of the β1 and β2 pro-peptides.

**Figure 4.**
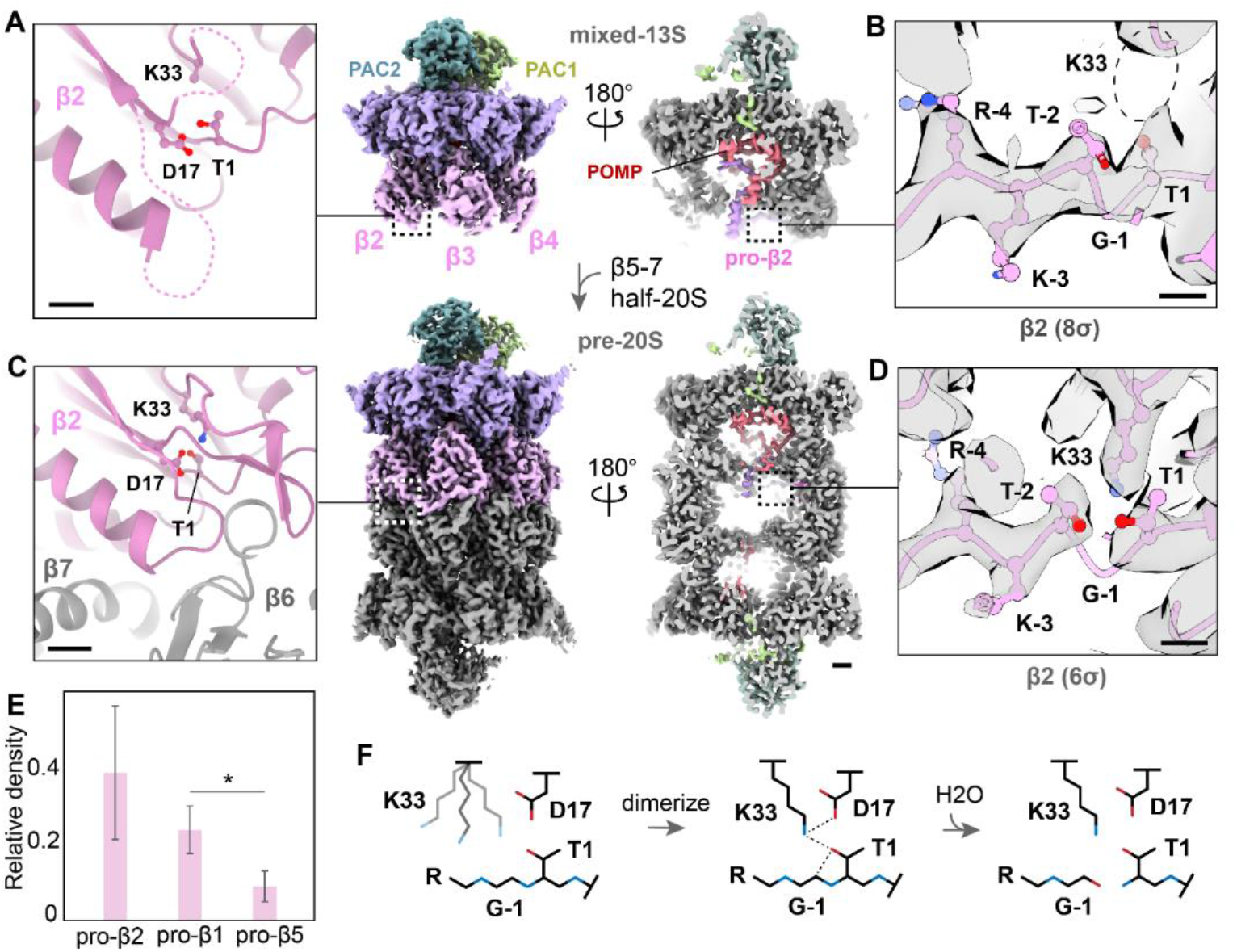
Pre-20S formation triggers autocatalysis of the β pro-peptides. (**A**) The cryo-EM map of mixed-13S shows poorly resolved density for the loops near the dimerization interface, indicating the region is likely flexible during β-ring assembly. Scale bar: 5 Å. (**B**) The sidechain of β2 K33 is not ordered in the mixed-13S structure, preventing cleavage of the β pro-peptide. Scale bar: 3 Å. (**C**) Dimerization of two half-proteasomes reorders the loops near the dimer interface into a stable structural arrangement. Scale bar: 5 Å. (**D**) Structural rearrangements induced by half-proteasome dimerization positions the sidechain of K33 into a catalytically active conformation. A break in the cryo-EM density (gray) can be observed between β2 T1 and T-2 at higher σ thresholds, indicating cleavage of the pro-peptide. Scale bar: 3 Å. (**E**) Quantification of the average cryo-EM map densities for the first six residues of the β pro-peptides after T1. *p-value<0.01 (**F**) Schematic depicting how structural rearrangement of K33 activates autocatalysis of the β pro-peptide. Scale bar for cryo-EM maps: 10 Å.

Each of the catalytic β-subunits contain a catalytic threonine T1, a conserved lysine K33, and a conserved aspartic acid D17 (*23, 47*–*49*). Cleavage occurs between T1 and the conserved G-

1 of the pro-peptide (Supplementary Figure 1C), which liberates the catalytic T1 residue of the β subunit. Focusing on β2, K33 and the adjacent loop region are disordered prior to half-proteasome dimerization (Figure 4A-B), disrupting the active site (*50*). Consistent with a catalytically inactive enzyme, strong density for the β2 pro-peptide is observed in the pre-13S and mixed-13S cryo-EM maps (Figure 4B, Supplementary Figure 8). Upon dimerization of two half-proteasomes, K33 and the adjacent loops of β2 are stabilized by interactions at the dimer interface (Figure 4C-D). This structural rearrangement places K33 into the correct position to coordinate T1 and D17, thus activating autocatalytic cleavage of the β2 pro-peptide. Consistent with an active enzyme, a break in the pre-20S cryo-EM map density between β2 T1 and G-1 of the pro-peptide can be observed at higher σ thresholds (Figure 4D, Supplementary Figure 8). Map density for the β2 pro-peptide can still be observed at higher σ thresholds, indicating that the pro-peptide can remain bound after cleavage.

### 20S maturation

The final steps of 20S assembly involve the removal of the β pro-peptides and dissociation of the bound chaperones, but how this occurs remains unclear. Examination of the pre-13S and mixed-13S cryo-EM maps show that the chaperones and β2 pro-peptide are bound stoichiometrically relative to β2 (Figure 5A-B, Supplementary Figure 9). In the pre-20S cryo-EM map, both the chaperones and β pro-peptides have lower densities compared to the α and β subunits, indicating that the pre-20S cryo-EM map likely represents a mixture of maturing 20S complexes. The β2 pro-peptide and the N-terminal region of POMP have substantially lower density in the pre-20S map. The C-terminal tail of PAC1 is also significantly lower in density relative to the rest of the protein in the pre-20S map. The relative differences in density observed within each component indicate that the final maturation of the pre-20S complex occurs through multiple stages. Because the proteins were prepared at 4°C, we reasoned that the complexes would mature further at physiological temperatures. Indeed, incubation of purified pre-20S complexes at 37°C for 1 h prior to cryo-EM grid preparation and analysis revealed a more mature complex, which we designate pre-20S’ (Figure 5B, Supplementary Figures 2D, 3, 9). The cryo-EM map of pre-20S’ shows lower density for the β2 pro-peptide, PAC1, PAC2, and POMP. Taken together, these results support a 3-stage model of 20S maturation (Figure 5C). First, the catalytic β subunits are activated through the cleavage and displacement of the β pro-peptides. Second, dissociation and degradation of the β2 pro-peptide causes displacement of the N-terminal region of POMP, which occurs concurrently with the destabilization of the PAC1 C-terminal tail. Finally, the C-terminal region of POMP dissociates and is degraded and PAC1/PAC2 is released to yield the mature 20S proteasome.

**Figure 5.**
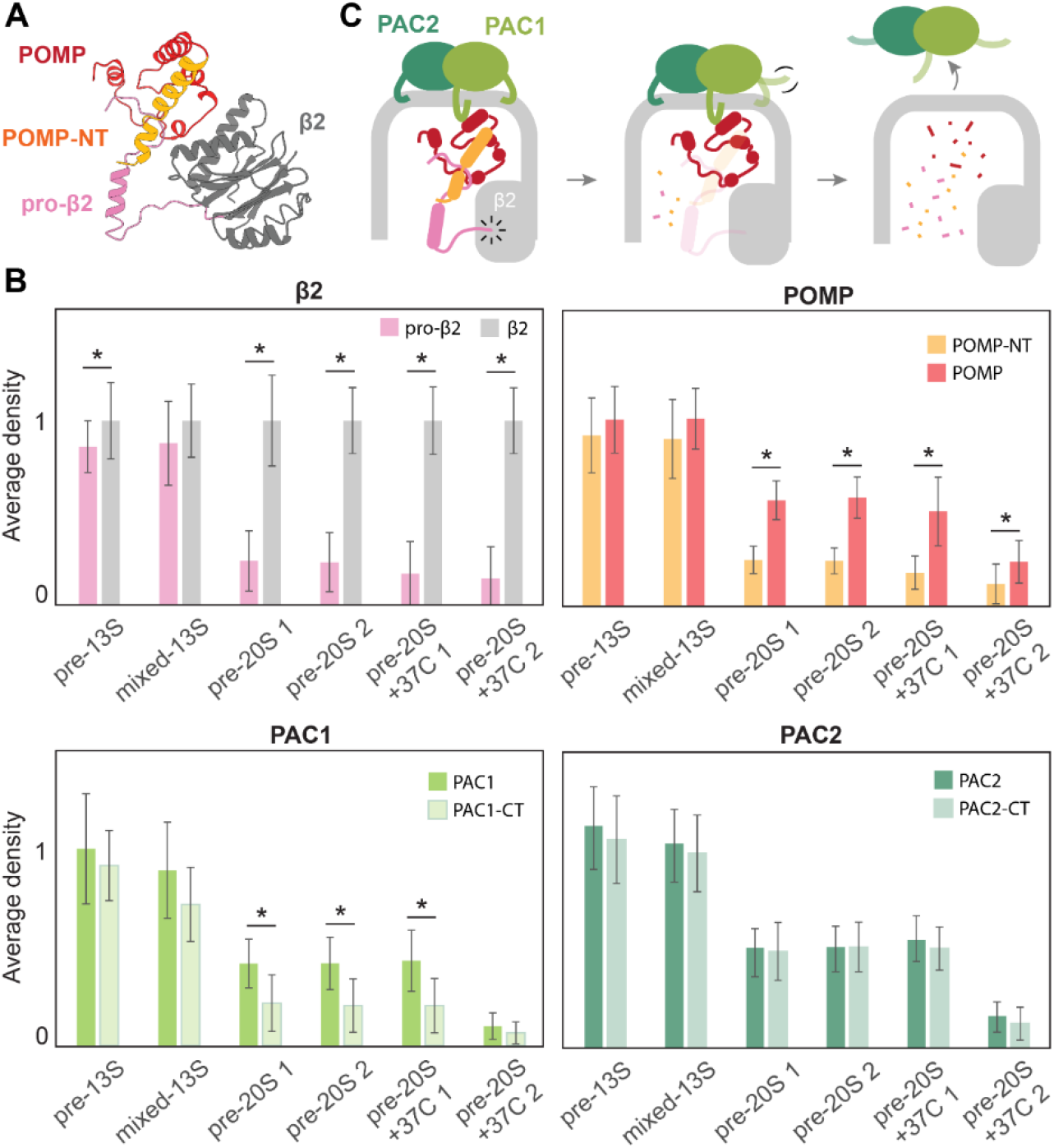
20S maturation proceeds in three stages. (**A**) Isolated view of the POMP interaction with β2 from the pre-13S atomic model. The β2 pro-peptide (pro-β2) interacts with the N-terminal region of POMP (POMP-NT). (**B**) Quantification of the cryo-EM density map values corresponding to β2, POMP, PAC1, and PAC2 shows a decrease in chaperone occupancy over the course of 20S maturation. Examination of specific regions within the proteins reveals a correlation between density changes in pro-β2, POMP-NT, and the C-terminal tail of PAC1 (PAC1-CT). Meanwhile, density changes in the C-terminal portion of POMP are correlated with changes in the N-terminal portion of PAC1 and PAC2. *p-value<0.001. (**C**) Cartoon depicting the 3-stage progression of 20S maturation involving β pro-peptide cleavage, pro-β2 and POMP-NT dissociation, and POMP/PAC1/PAC2 release and degradation.

## DISCUSSION

The tagging of endogenous proteins helps to preserve their native intracellular states and is a powerful approach for the study of complex biological processes. By tagging specific chaperones, we were able to capture molecular snapshots at each major stage of the 20S proteasome assembly pathway, allowing us to decipher the individual steps in the process. The structures reveal how PAC1-4 help to initiate assembly by binding to α4-7. While α1-3 do not interact meaningfully with PAC1-4, they make substantial contacts with POMP, suggesting that the binding of POMP may serve as a checkpoint to ensure the proper formation of the α-ring. The N terminus of POMP and the pro-peptide of β2 clash with PAC3/PAC4, which likely causes dissociation of the chaperones upon binding of β3. The clash between POMP/β2 and PAC3/PAC4 in our structures of the human proteins is substantially less severe than what has been predicted for the yeast homologs (*25*), suggesting a potential species-level difference in proteasome assembly at this stage. Dimerization of half-proteasomes stabilizes the critical lysine K33 and activates autocatalysis of the β pro-peptides, which likely occurs prior to the release of POMP and PAC1/PAC2. This is supported by the relatively lower map densities for the β pro-peptides in the pre-20S and pre-20S’ maps, as well as the fact that POMP needs to be degraded by active β subunits. The dissociation of the β2 pro-peptide and the associated N-terminal region of POMP likely destabilize the binding of POMP and PAC1/PAC2, thus promoting their release from the mature 20S complex.

One outstanding question concerns the role of the PAC1 N-terminal tail in 20S assembly, which inserts into the α-ring pore (*25*). It has been suggested that the PAC1 N-terminal tail may be involved in α-ring formation. However, the N-terminal tail of PAC1 is not inserted into the α-ring pore in the PAC1-4/α-ring structure from this study. Therefore, it is unlikely that the PAC1 N terminus plays a role in α-ring formation and its function remains unclear. The N terminus of PAC1 is inserted in the α-ring pore of the pre-13S, mixed-13S, and pre-20S structures, indicating a structural rearrangement after completion of the α-ring and initiation of β-ring formation. It is possible that insertion of the PAC1 N-terminal tail helps to stabilize the binding of PAC1/PAC2 to the correctly formed α-ring. The N-terminal tail of PAC1 also helps to block the open α-ring gate to prevent substrate entry into the immature proteasome (*25*). Additionally, the N terminus of PAC1 interacts with POMP, which may be forming a signal relay network that aids in the release of the chaperones from the maturing 20S complex.

The β5 pro-peptide is processed more quickly relative to the pro-peptides of β1 and β2, but the mechanistic basis for this is unclear. β5 contains a conserved histidine at the −2 position while threonine occupies the same position in β1 and β2 (Supplementary Figure 1C). The conserved histidine may be serving as a second proton acceptor in β5, which is a role that is fulfilled exclusively by K33 in β1 and β2. The two proton acceptors in β5 (K33 and H-2) may help to boost the autocatalysis rate of the pro-peptide. Indeed, mutation of H-2 to phenylalanine reduces but does not abrogate cleavage of the β5 pro-peptide (*45, 47*). In contrast, mutation of the conserved histidine to lysine, which also functions as a proton acceptor, preserves autocatalytic activity (*45, 47*). The rapid processing of the β5 pro-peptide may help to catalyze the degradation of other β subunit pro-peptides and POMP, which would help promote the maturation of the pre-20S complex into functional 20S proteasomes.

## METHODS

### Primers for gRNA Synthesis

ML557

TAATACGACTCACTATAG

ML558

AAAAAAAGCACCGACTCGGTGC

ML611

AAAAAAAGCACCGACTCGGTGCCACTTTTTCAAGTTGATAACGGACTAGCCTTATTTAAACTTGCTATGCTGTTTCCAGCATAGCTCTTAAAC

<u>PSMG2</u> oligo

TAATACGACTCACTATAG<u>GCGACCATGTTCGTT CCCTG</u>GTTTAAGAGCTATGCTGGAA

<u>PSMG3</u> oligo

TAATACGACTCACTATAG<u>CGCCATGGAAGACAC GCCGT</u>GTTTAAGAGCTATGCTGGAA

<u>POMP</u> oligo

TAATACGACTCACTATAG<u>GAGCTGCGGAAGAT GGTGAG</u>GTTTAAGAGCTATGCTGGAA

### Single-Strand DNA for Homology-Directed Repair (HDR)

PSMG2(β4)-nt-<u>mNG2(11)</u>-*<u>StrepII</u>*

TCTTGCCAGGGCCGCGGTTAGTCCCTGCTGGC CACCCCACTGCGACCATG<u>ACCGAGCTCAACTT CAAGGAGTGGCAAAAGGCCTTTACCGATATGA TG</u>*<u>TGGTCCCACCCTCAATTCGAGAAG</u> GGGAGT GCG*TTCGTTCCCTGCGGGGAGTCGGCCCCCG ACCTTGCCGGCTTCACCCTCCT

PSMG3(PAC3)-nt-<u>twinStrep</u>

TTTTTTCCCCTTTCTTTTTAACCTAAATTAAAGCT GCCACTGCAGAGCCCCGCCATG<u>TGGTCCCACC CTCAATTCGAGAAGGGAGGGGGGTCCGGTGGAGGTTCTGGGGGCTCTGCTTGGTCACATCCGC AGTTTGAGAAA</u>GAAGACACGCCGTTGGTGATAT CGAAGCAGAAGACGGAGGTGGTGTGCGGGGT CCCCA

POMP-nt-<u>mNG2(11)</u>-*<u>StrepII</u>*

GGCGGGGTCGACTGACGGTAACGGGGCAGAG AGGCTGTTCGCAGAGCTGCGGAAGATG<u>ACCGA GCTCAACTTCAAGGAGTGGCAAAAGGCCTTTA CCGATATGATG</u>*<u>TGGTCCCACCCTCAATTCGAGA AG</u>*GTGAGTGGGTACCCGGGCGGCTGGAGTTC CACGCGGGCTCGGGACCGTGGCTCGGCAGAAACAGGCAGTC

### Cas9 purification

The pET15-SpCas9-NLS-HIS plasmid was purchased from Addgene (#62731) (*51*). T7 Express cells (New England Biolabs) were transformed with the plasmid and grown in 5 ml of LB media (VWR) supplemented with 100 μg/ml carbenicillin (final concentration) overnight at 37 °C with shaking at 225 RPM. The next day, the cells were added to 100 ml of LB + carbenicillin (100 μg/ml) and incubated at 37 °C with shaking at 225 RPM. When OD600 reached 0.8, the 100 ml culture was added to 1 l of pre-warmed LB + carbenicillin (100 μg/ml) + 0.5% glucose and incubated at 37 °C with shaking at 225 RPM. When OD600 reached 0.4, the temperature was reduced to 16 °C. When OD600 reached 0.6, protein expression was induced with 0.2 mM IPTG (final concentration) overnight. The next day, cells were harvested by centrifugation at 4,000g for 10 mins and resuspended in 20 ml HBS buffer (50 mM HEPES, pH 7, 300 mM NaCl) supplemented with 25 mM imidazole, 100 ul HALT protease inhibitor cocktail (ThermoFisher), and 4 ml of 3 M NaCl. Protein purification was carried out at 4 °C. Cells were lysed by sonication and large debris were removed by centrifugation at 4,000g for 5 mins. The supernatant was centrifuged at ~100,000g for 20 mins in an ultracentrifuge (Beckman) to remove small insoluble aggregates. The supernatant was passed through a 0.22 μm syringe filter prior to adding to 0.5 ml NEBExpress Ni Resin (New England Biolabs) by gravity flow. The beads were washed with 3×1 ml of HBS buffer supplemented with 25 mM imidazole. Protein was eluted with 3×1 ml of HBS supplemented with 300 mM imidazole and loaded onto a 1 ml HiTrap SP column equilibrated with buffer A (50 mM HEPES, 400 mM NaCl, pH 7). The column was washed with 3 ml of buffer A and bound proteins were eluted with 3 ml of buffer B (50 mM HEPES, pH 7, 650 mM NaCl). The elution was diluted 3x with buffer C (50 mM HEPES, pH 7, 150 mM NaCl) and supplemented with 1 mM DTT and 10 % glycerol (final concentrations). Protein was concentrated in a 100 M.W. centrifugal device (Millipore) to ~13 mg/ml, aliquoted, and stored at −80 °C.

### Cas9 HDR Knock-Ins

Cas9 HDR knock-ins were performed as previously described (*31*) with slight modifications. The gRNA IVT template was generated by PCR in 20 μl reactions containing 8 μl Milli-Q water, 1 μl ML557+558 (50 uM stock), 0.5 μl ML611 (2 uM stock), 0.5 μl gene-specific oligo (2 uM stock), and 10 μl Q5 2X master mix. The PCR products were purified with the DNA clean & concentrator kit (Zymo Research, D4014) and eluted with 10 μl Milli-Q water. The gRNA was synthesized using the HiScribe T7 Quick High Yield RNA Synthesis Kit (New England Biolabs, E2050S) in 20 ul reactions containing 6 μl Milli-Q water, 1 μl DNA template (50-100 ng total), 10 μl NTP buffer mix, 2 μl T7 10X master mix, and 1 μl RNAsin (Promega). The reaction mixture was incubated at 37 °C for 16 h overnight and the gRNA was purified using the RNA clean & concentrator kit (Zymo Research, R1014). The gRNA was eluted in 10 μl Milli-Q water and 1 μl of 1.5 M NaCl (150 mM final concentration) was added to the RNA elution to prevent Cas9 precipitation during RNP reconstitution. The gRNA was incubated at 70 °C for 5 mins and stored at −20 °C until use. The Cas9-gRNA RNP was reconstituted by adding 20 ug gRNA (2-5 ul) to 5 μl Cas9 (80 uM; 13 mg/ml) at room temperature. 1.5 μl of ssDNA HDR template (IDT, 100 uM stock) was added to the Cas9-gRNA mixture. For each knock-in, 1×10^6 Expi293F cells were spun down at 500 g for 3 min and resuspended in 15 μl of Expi293 expression medium (gibco). 10 ul of cells were added to the Cas9 RNP and mixed well. The mixture was gently transferred to a Lonza Nucleofector X unit S 16-well strip without generating air bubbles. Program FS-100 was executed in a Lonza 4D Nucleofector and cells were immediately transferred to a 6-well plate containing 2 ml of Expi293 expression medium with 2.9 μl HDR enhancer (IDT). The plate was incubated at 37 °C, 8 % CO2 with shaking at 125 RPM. After 24 h, the media was replaced with fresh media without HDR enhancer.

### FACS for PSMG2(β4)-mNG2(11)-StrepII

For the PSMG2(PAC2)-mNG2(11)-StrepII knock-in, cells were sorted by fluorescence assisted cell sorting (FACS) to isolate cells with successful incorporation of the tag (Supplementary Figure 1D). When cells have recovered (7-10 days after knock-in), cells were transferred to 10 ml of Expi293 expression medium in a 125 ml flask and incubated at 37 °C, 8 % CO2, with shaking at 125 RPM. When cells reached ~2×10^6/ml, cells were diluted to 1×10^6/ml in 20 ml media. Cells were transfected with 20 ug of pcDNA5-mNG2(1-10) plasmid using Expifectamine 293 (gibco). 2 days after transfection, fluorescent cells containing successfully incorporated tags were sorted by FACS at the Sanford Burnham Prebys Flow Cytometry Shared Resource. Sorted cells were expanded, aliquoted, and stored in liquid nitrogen.

### Clonal Selection of knock-in cell lines

Clonal selection was carried out on PSMG3(PAC3)-twinstrep and POMP-mNG2(11)-StrepII knock-ins to isolate cells with successfully incorporated tags. Cells were seeded at ~100 cells/well in a 96-well plate and incubated at 37 °C, 8% CO2. After the cells grew to large colonies (>100 cells), individual colonies were extracted under a Leica DM IL inverted light microscope using a pipette and seeded into a new well with fresh Expi293 expression medium containing Penicillin-Streptomycin (Cytiva). The plate was incubated at 37 °C, 8% CO2. After 10-14 days, cells were transferred to 2 ml of fresh Expi293 expression medium in a 6-well plate and incubated at 37 °C, 8% CO2. When cells reached 1×10^6/ml, 1 ml of cells were spun down at 1000g and used for sequencing. Genomic DNA was extracted from cells using the Zymo Plasmid Miniprep kit (Zymo Research, D4209) and eluted in 25 μl Milli-Q water. DNA 100-700 bp upstream and downstream of the tag incorporation site was amplified by PCR. PCR products were purified using the DNA clean & concentrator kit (Zymo Research, D4014) and sequenced by Genewiz. Positive clones were expanded, aliquoted, and stored in liquid nitrogen.

### Cell Culture and Proteasome Purification

2×100 ml cultures of each cell line were grown in Expi293 expression medium (gibco) at 37 °C, 8% CO_2_, and shaking at 125 RPM. When cells reached a density of 3-4×10^6/ml, cells were harvested by centrifugation at 500g for 5 min. Cells were resuspended in 10 ml HBS buffer (50 mM Hepes, 150 mM NaCl, pH 7.2) with the addition of DTT (1 mM final concentration), 100 ul of 100X HALT protease inhibitor cocktail (ThermoFisher), and 200 ul BioLock (iba). Cells were lysed by sonication and insoluble debris was removed by centrifugation at 4,000g for 10 mins. The supernatant was centrifuged at ~100,000g for 20 min at 4°C in an ultracentrifuge. The supernatant was filtered through a 0.2 μm syringe filter and transferred to a column containing 50 μl Strep-Tactin XT Sepharose resin (Cytiva) by gravity flow. The resin was washed with 4×500 μl HBS buffer and the bound proteins were eluted with 1 ml HBS buffer with 50 mM biotin (final concentration). The eluted proteins were concentrated in a 50,000 MW centrifugal concentrator (Millipore).

### Cryo-EM Imaging and Data Processing

For PSMG2(PAC2)-tagged samples, 2.5 μl of purified protein (1.3mg/ml) were applied to Quantifoil Cu 300 mesh R 2/2 grids (glow-discharged in a Pelco easiGlow at 15 mA for 25s), blotted for 10 s (4 °C, 100 % humidity, blot force 0) using Whatman 1 filter paper, and plunge frozen in liquid ethane using a Vitrobot Mark IV (ThermoFisher). For PSMG3(PAC3) and POMP-tagged samples, 2.5 μl of purified protein (0.5 mg/ml) was applied to Quantifoil Cu 300 mesh R 2/2 grids coated with graphene oxide (Sigma) (*52*), blotted for 1 s (4 °C, 100 % humidity, blot force 0) using Whatman 1 filter paper, and plunge frozen in liquid ethane using a Vitrobot Mark IV (ThermoFisher). The vitrified samples were imaged in a Titan Krios electron microscope (FEI/ThermoFisher) in the Cryo-EM facility of the Structural Biology Shared Resource at Sanford Burnham Prebys Medical Discovery Institute. Images were collected on a Gatan K3 detector using SerialEM (*53*) under super-resolution settings (*54*). (Extended Data Table 1). Image processing of the micrographs were carried out in cryoSPARC (*55, 56*), including motion correction, CTF estimation, particle picking, classification, and refinement. Resolution of the refined cryo-EM maps were estimated by gold-standard Fourier Shell Correlation at the 0.143 criterion (*57*). Directional FSC of the maps were calculated as described previously (*58*). Atomic models were built into the cryo-EM density maps using Coot (*59, 60*) and Phenix (*61, 62*). Initial models used for model building were Protein Data Bank (PDB) ID codes 7NAN (20S), 6JPT (PAC3), and 5WTQ (PAC4). The resulting cryo-EM maps and models were visualized using UCSF Chimera and UCSF ChimeraX (*63, 64*).

### Density analysis and statistics

The average cryo-EM map density values corresponding to individual subunits and chaperones were calculated per residue of an atomic model using the ‘Values at Atom Positions’ function in Chimera and analyzed in Microsoft Excel. Only backbone atoms were considered in the analysis. Statistical significance was evaluated using a two-tailed t-test with unequal variance to calculate p-values.

## DATA AVAILABILITY

The atomic models and cryo-EM maps were deposited in the Protein Data Bank and Electron Microscopy Data Bank under accession codes 8TM3 and 41377 (PAC1-4/α-ring), 8TM4 and 41378 (pre-13S), 8TM5 and 41379 (mixed-13S), 8TM6 and 41380 (pre-20S), and 41381 (pre-20S’).

## ACKNOWLEDGEMENTS

We would like to thank Laura Koepping from the Sanford Burnham Prebys Medical Discovery Institute (SBP) Cryo-EM facility of the Structural Biology Shared Resource and Amy Cortez from the SBP Flow Cytometry Shared Resource for their amazing support in the cryo-EM and cell sorting work, respectively. We would also like to thank Yifan Cheng (University of California San Francisco) and Elena Pasquale (SBP) for critical reading of the manuscript.

This work is supported by National Institutes of Health (NIH) grant R35 GM147487 to JZ. This work is also supported by National Cancer Institute Cancer Center Support Grant P30 CA030199 and NIH instrumentation grant S10 OD026926. Molecular graphics and analyses performed with UCSF ChimeraX, developed by the Resource for Biocomputing, Visualization, and Informatics at the University of California, San Francisco, with support from NIH R01-GM129325 and the Office of Cyber Infrastructure and Computational Biology, National Institute of Allergy and Infectious Diseases.

## AUTHOR CONTRIBUTIONS

H.Z. and J.Z. conceived the project and designed the experiments. H.Z. and C.Z. generated and maintained the cell lines. H.Z. and Z.M. purified the proteins and prepared the samples. H.Z. collected and processed the data. H.Z. and J.Z. analyzed the data. H.Z., C.Z., and J.Z. prepared the figures and wrote the manuscript.

## REFERENCES

1. A. Rousseau, A. Bertolotti, Regulation of proteasome assembly and activity in health and disease. Nat. Rev. Mol. Cell Biol. 2018 1911. 19, 697–712 (2018).

2. Y. Xie, A. Varshavsky, RPN4 is a ligand, substrate, and transcriptional regulator of the 26S proteasome: A negative feedback circuit. Proc. Natl. Acad. Sci. U. S. A. 98, 3056–3061 (2001).

3. M. Ma, Z. L. Liu, Comparative transcriptome profiling analyses during the lag phase uncover YAP1, PDR1, PDR3, RPN4, and HSF1 as key regulatory genes in genomic adaptation to the lignocellulose derived inhibitor HMF for Saccharomyces cerevisiae. BMC Genomics. 11, 1–19 (2010).

4. S. Meiners, D. Heyken, A. Weller, A. Ludwig, K. Stangl, P. M. Kloetzel, E. Krüger, Inhibition of Proteasome Activity Induces Concerted Expression of Proteasome Genes and de Novo Formation of Mammalian Proteasomes. J. Biol. Chem. 278, 21517–21525 (2003).

5. M.-K. Kwak, N. Wakabayashi, J. L. Greenlaw, M. Yamamoto, T. W. Kensler, Antioxidants Enhance Mammalian Proteasome Expression through the Keap1-Nrf2 Signaling Pathway. Mol. Cell. Biol. 23, 8786–8794 (2003).

6. S. K. Radhakrishnan, C. S. Lee, P. Young, A. Beskow, J. Y. Chan, R. J. Deshaies, Transcription Factor Nrf1 Mediates the Proteasome Recovery Pathway after Proteasome Inhibition in Mammalian Cells. Mol. Cell. 38, 17–28 (2010).

7. A. Rousseau, A. Bertolotti, An evolutionarily conserved pathway controls proteasome homeostasis. Nat. 2016 5367615. 536, 184–189 (2016).

8. K. L. Rock, C. Gramm, L. Rothstein, K. Clark, R. Stein, L. Dick, D. Hwang, A. L. Goldberg, Inhibitors of the proteasome block the degradation of most cell proteins and the generation of peptides presented on MHC class I molecules. Cell. 78, 761–771 (1994).

9. K. A. Opoku-Nsiah, J. E. Gestwicki, Aim for the core: suitability of the ubiquitin-independent 20S proteasome as a drug target in neurodegeneration. Transl. Res. 198, 48–57 (2018).

10. S. H. Jung, S. H. Jae, I. Chang, S. Kim, Age-Associated Decrease in Proteasome Content and Activities in Human Dermal Fibroblasts: Restoration of Normal Level of Proteasome Subunits Reduces Aging Markers in Fibroblasts From Elderly Persons. Journals Gerontol. Ser. A. 62, 490–499 (2007).

11. A. Ciechanover, Y. T. Kwon, Degradation of misfolded proteins in neurodegenerative diseases: therapeutic targets and strategies. Exp. Mol. Med. 2015 473. 47, e147–e147 (2015).

12. D. Vilchez, I. Saez, A. Dillin, The role of protein clearance mechanisms in organismal ageing and age-related diseases. Nat. Commun. 5 (2014), doi:10.1038/ncomms6659.

13. P. Walter, D. Ron, The unfolded protein response: From stress pathway to homeostatic regulation. Science (80-.). 334, 1081–1086 (2011).

14. A. Hanssum, Z. Zhong, A. Rousseau, A. Krzyzosiak, A. Sigurdardottir, A. Bertolotti, An Inducible Chaperone Adapts Proteasome Assembly to Stress. Mol. Cell. 55, 566–577 (2014).

15. Y. Hirano, K. B. Hendil, H. Yashiroda, S. Iemura, R. Nagane, Y. Hioki, T. Natsume, K. Tanaka, S. Murata, A heterodimeric complex that promotes the assembly of mammalian 20S proteasomes. Nat. 2005 4377063. 437, 1381–1385 (2005).

16. Y. Hirano, H. Hayashi, S. Iemura, K. B. Hendil, S. Niwa, T. Kishimoto, M. Kasahara, T. Natsume, K. Tanaka, S. Murata, Cooperation of Multiple Chaperones Required for the Assembly of Mammalian 20S Proteasomes. Mol. Cell. 24, 977–984 (2006).

17. B. Le Tallec, M.-B. Barrault, R. Courbeyrette, R. Guérois, M.-C. Marsolier-Kergoat, A. Peyroche, 20S Proteasome Assembly Is Orchestrated by Two Distinct Pairs of Chaperones in Yeast and in Mammals. Mol. Cell. 27, 660–674 (2007).

18. P. C. Ramos, J. Höckendorff, E. S. Johnson, A. Varshavsky, R. J. Dohmen, Ump1p Is Required for Proper Maturation of the 20S Proteasome and Becomes Its Substrate upon Completion of the Assembly. Cell. 92, 489–499 (1998).

19. L. Burri, J. Höckendorff, U. Boehm, T. Klamp, R. J. Dohmen, F. Lévy, Identification and characterization of a mammalian protein interacting with 20S proteasome precursors. Proc. Natl. Acad. Sci. U. S. A. 97, 10348–10353 (2000).

20. E. Witt, D. Zantopf, M. Schmidt, R. Kraft, P. M. Kloetzel, E. Krüger, Characterisation of the newly identified human Ump1 homologue POMP and analysis of LMP7(β5i) incorporation into 20 S proteasomes. J. Mol. Biol. 301 (2000), doi:10.1006/jmbi.2000.3959.

21. T. A. Griffin, J. P. Slack, T. S. McCluskey, J. J. Monaco, R. A. Colbert, Identification of proteassemblin, a mammalian homologue of the yeast protein, Ump1p, that is required for normal proteasome assembly. Mol. Cell Biol. Res. Commun. 3 (2000), doi:10.1006/mcbr.2000.0213.

22. E. Seemuller, A. Lupas, W. Baumeister, Autocatalytic processing of the 20S proteasome. Nat. 1996 3826590. 382, 468–470 (1996).

23. G. Schmidtke, R. Kraft, S. Kostka, P. Henklein, C. Frömmel, J. Löwe, R. Huber, P. M. Kloetzel, M. Schmidt, Analysis of mammalian 20S proteasome biogenesis: the maturation of beta-subunits is an ordered two-step mechanism involving autocatalysis. EMBO J. 15, 6887–6898 (1996).

24. Y. Hirano, T. Kaneko, K. Okamoto, M. Bai, H. Yashiroda, K. Furuyama, K. Kato, K. Tanaka, S. Murata, Dissecting β-ring assembly pathway of the mammalian 20S proteasome. EMBO J. 27, 2204–2213 (2008).

25. H. M. Schnell, R. M. Walsh, S. Rawson, M. Kaur, M. K. Bhanu, G. Tian, M. A. Prado, A. Guerra-Moreno, J. A. Paulo, S. P. Gygi, J. Roelofs, D. Finley, J. Hanna, Structures of chaperone-associated assembly intermediates reveal coordinated mechanisms of proteasome biogenesis. Nat. Struct. Mol. Biol. 28, 418–425 (2021).

26. M. Kock, M. M. Nunes, M. Hemann, S. Kube, R. Jürgen Dohmen, F. Herzog, P. C. Ramos, P. Wendler, Proteasome assembly from 15S precursors involves major conformational changes and recycling of the Pba1–Pba2 chaperone. Nat. Commun. 2015 61. 6, 1–10 (2015).

27. A. R. Kusmierczyk, M. J. Kunjappu, M. Funakoshi, M. Hochstrasser, A multimeric assembly factor controls the formation of alternative 20S proteasomes. Nat. Struct. Mol. Biol. 15, 237–244 (2008).

28. H. Yashiroda, T. Mizushima, K. Okamoto, T. Kameyama, H. Hayashi, T. Kishimoto, S. Niwa, M. Kasahara, E. Kurimoto, E. Sakata, K. Takagi, A. Suzuki, Y. Hirano, S. Murata, K. Kato, T. Yamane, K. Tanaka, Crystal structure of a chaperone complex that contributes to the assembly of yeast 20S proteasomes. Nat. Struct. Mol. Biol. 15, 228–236 (2008).

29. W. L. H. Gerards, J. Enzlin, M. Häner, I. L. A. M. Hendriks, U. Aebi, H. Bloemendal, W. Boelens, The human α-type proteasomal subunit HsC8 forms a double ringlike structure, but does not assemble into proteasome-like particles with the β-type subunits HsDelta or HsBPROS26. J. Biol. Chem. 272 (1997), doi:10.1074/jbc.272.15.10080.

30. I. Velichutina, P. L. Connerly, C. S. Arendt, X. Li, M. Hochstrasser, Plasticity in eucaryotic 20S proteasome ring assembly revealed by a subunit deletion in yeast. EMBO J. 23 (2004), doi:10.1038/sj.emboj.7600059.

31. J. Zhao, S. Makhija, C. Zhou, H. Zhang, Y. Q. Wang, M. Muralidharan, B. Huang, Y. Cheng, Structural insights into the human PA28–20S proteasome enabled by efficient tagging and purification of endogenous proteins. Proc. Natl. Acad. Sci. U. S. A. 119, e2207200119 (2022).

32. A. V. Anzalone, L. W. Koblan, D. R. Liu, Genome editing with CRISPR–Cas nucleases, base editors, transposases and prime editors. Nat. Biotechnol. 2020 387. 38, 824–844 (2020).

33. M. D. Leonetti, S. Sekine, D. Kamiyama, J. S. Weissman, B. Huang, A scalable strategy for high-throughput GFP tagging of endogenous human proteins. Proc. Natl. Acad. Sci. 113, E3501–E3508 (2016).

34. P. Zwickl, J. Kleinz, W. Baumeister, Critical elements in proteasome assembly. Nat. Struct. Biol. 1 (1994), doi:10.1038/nsb1194-765.

35. B. M. Stadtmueller, E. Kish-Trier, K. Ferrell, C. N. Petersen, H. Robinson, D. G. Myszka, D. M. Eckert, T. Formosa, C. P. Hill, Structure of a Proteasome Pba1-Pba2 Complex. J. Biol. Chem. 287, 37371–37382 (2012).

36. T. Satoh, M. Yagi-Utsumi, K. Okamoto, E. Kurimoto, K. Tanaka, K. Kato, Molecular and Structural Basis of the Proteasome α Subunit Assembly Mechanism Mediated by the Proteasome-Assembling Chaperone PAC3-PAC4 Heterodimer. Int. J. Mol. Sci. 2019, Vol. 20, Page 2231. 20, 2231 (2019).

37. K. Takagi, Y. Saeki, H. Yashiroda, H. Yagi, A. Kaiho, S. Murata, T. Yamane, K. Tanaka, T. Mizushima, K. Kato, Pba3-Pba4 heterodimer acts as a molecular matchmaker in proteasome α-ring formation. Biochem. Biophys. Res. Commun. 450 (2014), doi:10.1016/j.bbrc.2014.06.119.

38. D. M. Smith, S. C. Chang, S. Park, D. Finley, Y. Cheng, A. L. Goldberg, Docking of the Proteasomal ATPases’ Carboxyl Termini in the 20S Proteasome’s α Ring Opens the Gate for Substrate Entry. Mol. Cell. 27 (2007), doi:10.1016/j.molcel.2007.06.033.

39. A. R. Kusmierczyk, M. J. Kunjappu, R. Y. Kim, M. Hochstrasser, A conserved 20S proteasome assembly factor requires a C-terminal HbYX motif for proteasomal precursor binding. Nat. Struct. Mol. Biol. 2011 185. 18, 622–629 (2011).

40. W. Wu, K. Sahara, S. Hirayama, X. Zhao, A. Watanabe, J. Hamazaki, H. Yashiroda, S. Murata, PAC1-PAC2 proteasome assembly chaperone retains the core α4–α7 assembly intermediates in the cytoplasm. Genes to Cells. 23, 839–848 (2018).

41. H. M. Schnell, J. Ang, S. Rawson, R. M. Walsh, Y. Micoogullari, J. Hanna, Mechanism of proteasome gate modulation by assembly chaperones Pba1 and Pba2. J. Biol. Chem. 298, 101906 (2022).

42. S. Frentzel, B. Pesold-Hurt, A. Seelig, P. M. Kloetzel, 20 S proteasomes are assembled via distinct precursor complexes processing of LMP2 and LMP7 proproteins takes place in 13-16 S preproteasome complexes. J. Mol. Biol. 236 (1994), doi:10.1016/0022-2836(94)90003-5.

43. X. Li, A. R. Kusmierczyk, P. Wong, A. Emili, M. Hochstrasser, β-Subunit appendages promote 20S proteasome assembly by overcoming an Ump1-dependent checkpoint. EMBO J. 26, 2339–2349 (2007).

44. A. J. Marques, C. Glanemann, P. C. Ramos, R. J. Dohmen, The C-terminal Extension of the β7 Subunit and Activator Complexes Stabilize Nascent 20 S Proteasomes and Promote Their Maturation. J. Biol. Chem. 282, 34869–34876 (2007).

45. X. Li, Y. Li, C. S. Arendt, M. Hochstrasser, Distinct Elements in the Proteasomal β5 Subunit Propeptide Required for Autocatalytic Processing and Proteasome Assembly. J. Biol. Chem. 291, 1991–2003 (2016).

46. P. Chen, M. Hochstrasser, Autocatalytic Subunit Processing Couples Active Site Formation in the 20S Proteasome to Completion of Assembly. Cell. 86, 961–972 (1996).

47. E. M. Huber, W. Heinemeyer, X. Li, C. S. Arendt, M. Hochstrasser, M. Groll, A unified mechanism for proteolysis and autocatalytic activation in the 20S proteasome. Nat. Commun. 2016 71. 7, 1–10 (2016).

48. C. S. Arendt, M. Hochstrasser, Identification of the yeast 20S proteasome catalytic centers and subunit interactions required for active-site formation. Proc. Natl. Acad. Sci. U. S. A. 94 (1997), doi:10.1073/pnas.94.14.7156.

49. L. Ditzel, R. Huber, K. Mann, W. Heinemeyer, D. H. Wolf, M. Groll, Conformational constraints for protein self-cleavage in the proteasome. J. Mol. Biol. 279 (1998), doi:10.1006/jmbi.1998.1818.

50. S. Witt, Y. Do Kwon, M. Sharon, K. Felderer, M. Beuttler, C. V. Robinson, W. Baumeister, B. K. Jap, Proteasome Assembly Triggers a Switch Required for Active-Site Maturation. Structure. 14, 1179–1188 (2006).

51. D. S. D’Astolfo, R. J. Pagliero, A. Pras, W. R. Karthaus, H. Clevers, V. Prasad, R. J. Lebbink, H. Rehmann, N. Geijsen, Efficient intracellular delivery of native proteins. Cell. 161, 674–690 (2015).

52. R. S. Pantelic, J. C. Meyer, U. Kaiser, W. Baumeister, J. M. Plitzko, Graphene oxide: A substrate for optimizing preparations of frozen-hydrated samples. J. Struct. Biol. 170, 152–156 (2010).

53. D. N. Mastronarde, Automated electron microscope tomography using robust prediction of specimen movements. J. Struct. Biol. 152, 36–51 (2005).

54. X. Li, P. Mooney, S. Zheng, C. R. Booth, M. B. Braunfeld, S. Gubbens, D. A. Agard, Y. Cheng, Electron counting and beam-induced motion correction enable near-atomic-resolution single-particle cryo-EM. Nat Methods. 10, 584–590 (2013).

55. A. Punjani, J. L. Rubinstein, D. J. Fleet, M. A. Brubaker, CryoSPARC: Algorithms for rapid unsupervised cryo-EM structure determination. Nat. Methods. 14, 290–296 (2017).

56. J. Zivanov, T. Nakane, S. H. W. Scheres, Estimation of high-order aberrations and anisotropic magnification from cryo-EM data sets in RELION-3.1. urn:issn:2052-2525. 7, 253–267 (2020).

57. P. B. Rosenthal, R. Henderson, Optimal determination of particle orientation, absolute hand, and contrast loss in single-particle electron cryomicroscopy. J Mol Biol. 333, 721–745 (2003).

58. S. Dang, S. Feng, J. Tien, C. J. Peters, D. Bulkley, M. Lolicato, J. Zhao, K. Zuberbühler, W. Ye, L. Qi, T. Chen, C. S. Craik, Y. N. Jan, D. L. Minor, Y. Cheng, L. Y. Jan, Cryo-EM structures of the TMEM16A calciumactivated chloride channel. Nature (2017), doi:10.1038/nature25024.

59. P. Emsley, K. Cowtan, Coot: Model-building tools for molecular graphics. Acta Crystallogr. Sect. D Biol. Crystallogr. 60, 2126–2132 (2004).

60. P. Emsley, B. Lohkamp, W. G. Scott, K. Cowtan, Features and development of Coot. Acta Crystallogr. Sect. D Biol. Crystallogr. 66, 486–501 (2010).

61. P. D. Adams, P. V. Afonine, G. Bunkóczi, V. B. Chen, I. W. Davis, N. Echols, J. J. Headd, L. W. Hung, G. J. Kapral, R. W. Grosse-Kunstleve, A. J. McCoy, N. W. Moriarty, R. Oeffner, R. J. Read, D. C. Richardson, J. S. Richardson, T. C. Terwilliger, P. H. Zwart, PHENIX: A comprehensive Python-based system for macromolecular structure solution. Acta Crystallogr. Sect. D Biol. Crystallogr. 66, 213–221 (2010).

62. P. D. Adams, P. V. Afonine, G. Bunkóczi, V. B. Chen, N. Echols, J. J. Headd, L. W. Hung, S. Jain, G. J. Kapral, R. W. Grosse Kunstleve, A. J. McCoy, N. W. Moriarty, R. D. Oeffner, R. J. Read, D. C. Richardson, J. S. Richardson, T. C. Terwilliger, P. H. Zwart, The Phenix software for automated determination of macromolecular structures. Methods. 55, 94–106 (2011).

63. E. F. Pettersen, T. D. Goddard, C. C. Huang, G. S. Couch, D. M. Greenblatt, E. C. Meng, T. E. Ferrin, UCSF Chimera--a visualization system for exploratory research and analysis. J Comput Chem. 25, 1605–1612 (2004).

64. E. F. Pettersen, T. D. Goddard, C. C. Huang, E. C. Meng, G. S. Couch, T. I. Croll, J. H. Morris, T. E. Ferrin, UCSF ChimeraX: Structure visualization for researchers, educators, and developers. Protein Sci. 30, 70–82 (2021).

