## Supplementary for "Structural basis of human 20S proteasome biogenesis": supplementary_figures.pdf

**A**

α-ring

| gene | protein |
|---|---|
| PSMA6 | α1 |
| PSMA2 | α2 |
| PSMA4 | α3 |
| PSMA7 | α4 |
| PSMA5 | α5 |
| PSMA1 | α6 |
| PSMA3 | α7 |

β-ring

| gene | protein |
|---|---|
| PSMB6 | β1 |
| PSMB7 | β2 |
| PSMB3 | β3 |
| PSMB2 | β4 |
| PSMB5 | β5 |
| PSMB1 | β6 |
| PSMB4 | β7 |

chaperones

| gene | protein |
|---|---|
| PSMG1 | PAC1 |
| PSMG2* | PAC2 |
| PSMG3* | PAC3 |
| PSMG4 | PAC4 |
| POMP* | POMP |

*tagged

**B**

proteasomal complexes

| complex | components |
|---|---|
| PAC1-4/α1-7 | PAC1-4, α1-7 |
| pre-13S | PAC1-2, POMP, α1-7, β2-3 |
| 13S | PAC1-2, POMP, α1-7, β2-4 |
| mixed-13S | PAC1-2, POMP, α1-7, β2-5 |
| 15S | PAC1-2, POMP, α1-7, β1-6 |
| pre-20S | (PAC1-2, POMP, α1-7, β1-7) x 2 |
| 20S | (α1-7, β1-7) x 2 |

**C**

-30 -20 -10 1 17 33

β1 MAATLLAARGAGPAPAWGPEAFTPDWESREVSTGTTIMAVQFDGGVVLGADSRTTTGSYIANRVTDKLTP...

β1 pro-peptide

-40 -30 -20 -10 1 17 33

β2 MAAVSVYAPPVGGFSFDNCRRNAVLEADFAKRGYKLPKVRKTGTTIAGVVYKDGIVLGADTRATEGMVVADKNCSKIHF...

β2 pro-peptide

-50 -40 -30 -20 -10 1 17 33

β5 MALASVLERPLPVNQRGFFGLGGRADLLDLGPGSLSDGLSLAAPGWGVPEEPGIEMLHGTTTLAFKFRHGVIVAADSRATAGAYIASQTVKKVIE...

β5 pro-peptide

**D**

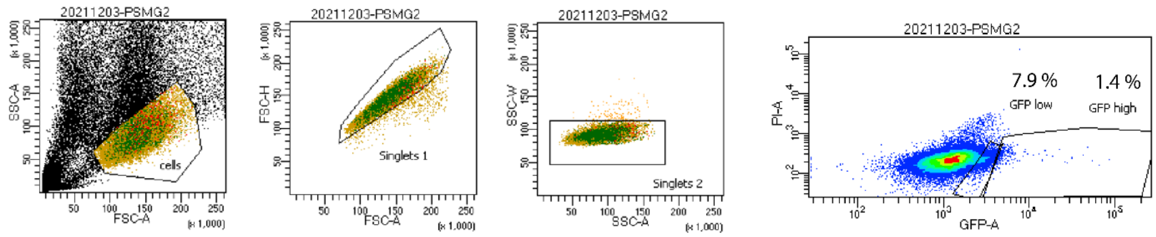

**Supplementary Figure 1. 20S proteasome components and FACS.** (**A**) Names of genes and corresponding proteins. Chaperones tagged in this study are indicated by '*'. (**B**) Names of proteasomal assembly intermediates and their corresponding components. (**C**) N-terminal amino acid sequences of β1, β2, and β5 showing their respective pro-peptides and conserved residues T1, D17, and K33. (**D**) Fluorescence cell sorting of PSMG2-mNG211_StrepII knock-in cells.

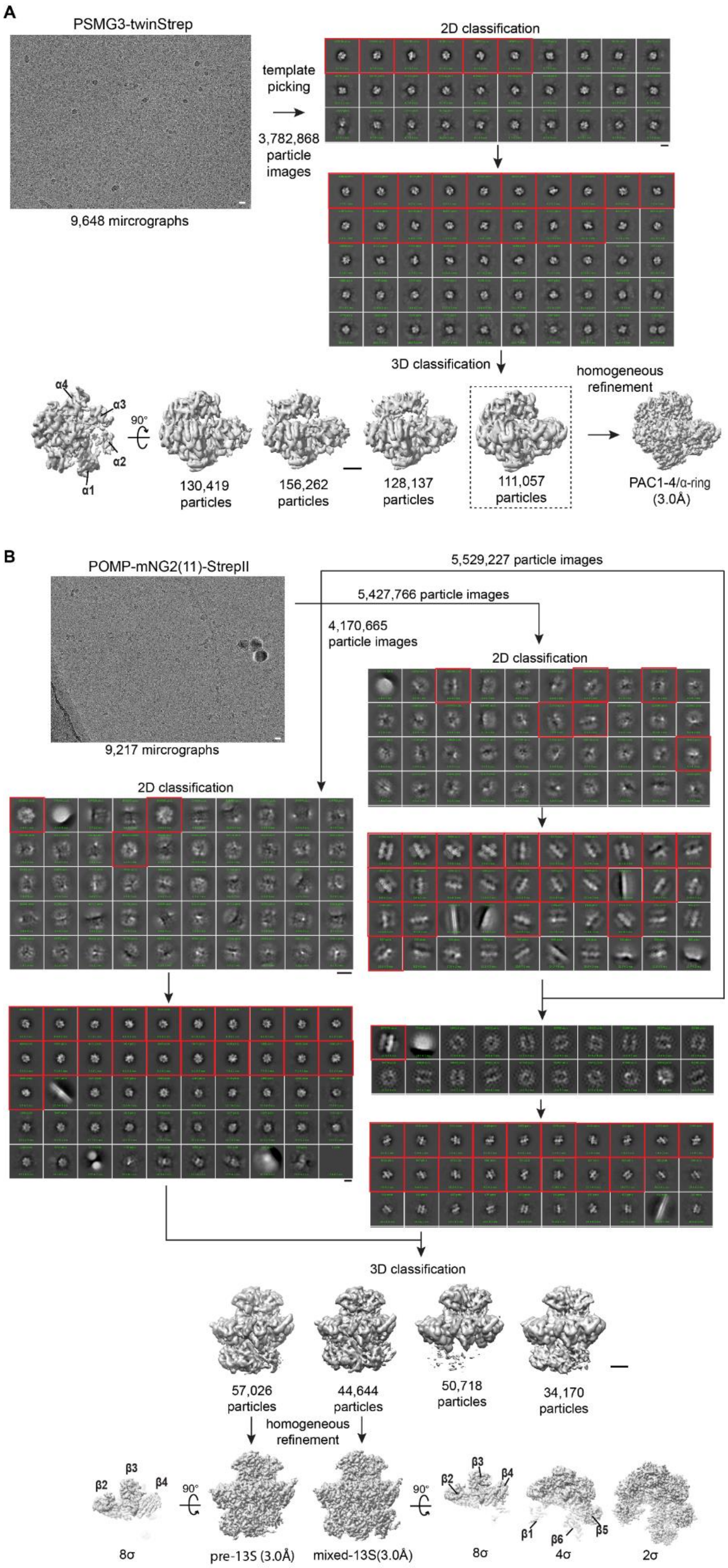
A
PSMG3-twinStrep
9,648 mircrographs
template picking
3,782,868 particle images
2D classification
3D classification
homogeneous refinement
α4
α3
α2
α1
90°
130,419 particles
156,262 particles
128,137 particles
111,057 particles
PAC1-4/α-ring (3.0Å)
B
POMP-mNG2(11)-StrepII
9,217 mircrographs
5,529,227 particle images
5,427,766 particle images
4,170,665 particle images
2D classification
2D classification
3D classification
57,026 particles
44,644 particles
50,718 particles
34,170 particles
homogeneous refinement
β2
β3
β4
90°
8σ
pre-13S (3.0Å)
mixed-13S(3.0Å)
90°
β2
β3
β4
β1
β6
β5
8σ
4σ
2σ

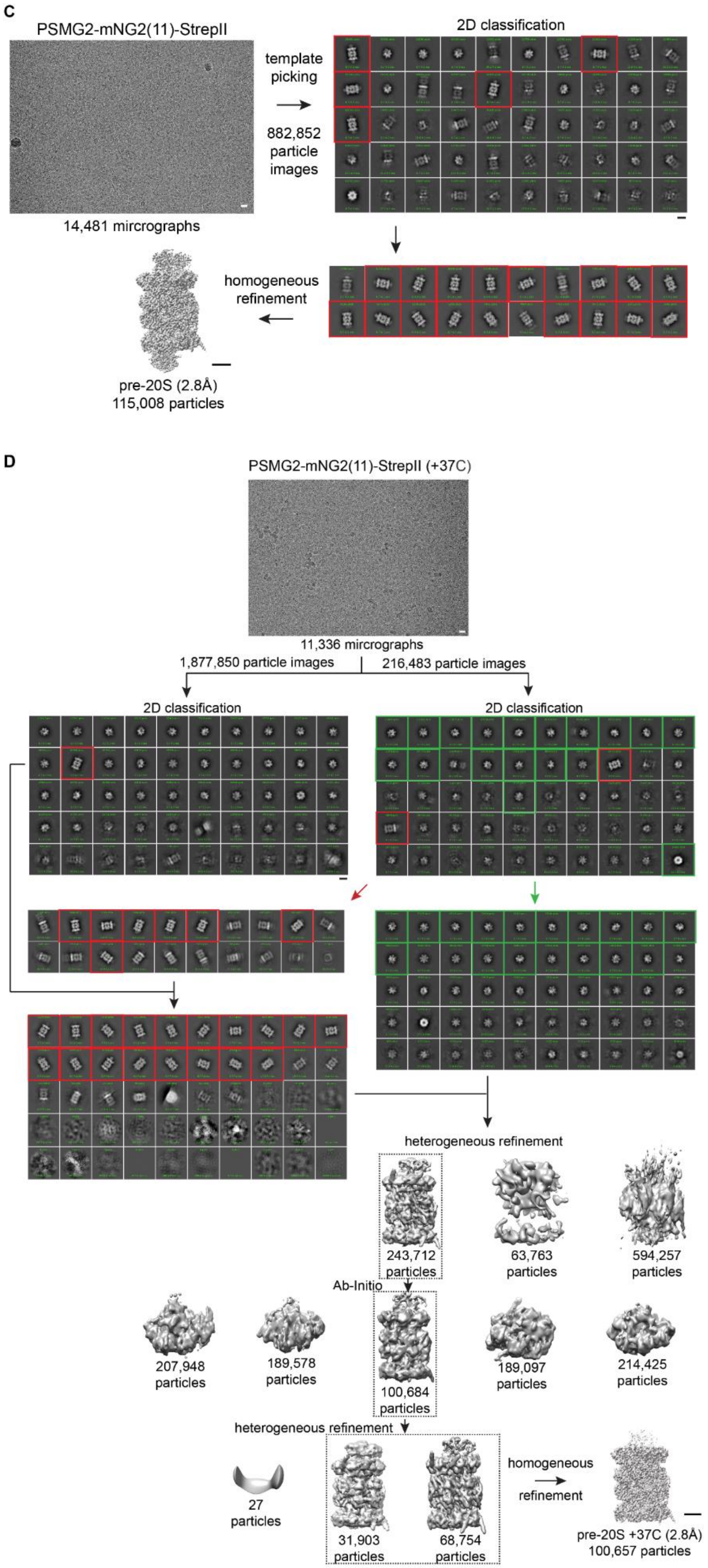

C
PSMG2-mNG2(11)-StrepII
template picking
882,852 particle images
14,481 mircrographs
2D classification
homogeneous refinement
pre-20S (2.8Å)
115,008 particles
D
PSMG2-mNG2(11)-StrepII (+37C)
11,336 mircrographs
1,877,850 particle images
216,483 particle images
2D classification
2D classification
heterogeneous refinement
243,712 particles
63,763 particles
594,257 particles
Ab-Initio
207,948 particles
189,578 particles
100,684 particles
189,097 particles
214,425 particles
heterogeneous refinement
27 particles
31,903 particles
68,754 particles
homogeneous refinement
pre-20S +37C (2.8Å)
100,657 particles

**Supplementary Figure 2. Cryo-EM image processing workflow.** (**A**) Analysis of PSMG3-twinStrep samples resulted in cryo-EM maps of PAC1-4/α-ring. 3D classification separated structures with stoichiometric and sub-stoichiometric α-rings. Scale bars: micrograph, 200 Å; 2D classes, 100 Å; cryo-EM maps, 20 Å. (**B**) Analysis of POMP-mNG2(11)-StrepII samples resulted in cryo-EM maps of pre-13S and mixed-13S. Scale bars: micrograph, 200 Å; 2D classes, 25 Å; cryo-EM maps, 20 Å. (**C**) Analysis of PSMG2-mNG2(11)-StrepII samples resulted in a cryo-EM map of pre-20S. Scale bars: micrograph, 200 Å; 2D classes, 100 Å; cryo-EM map, 20 Å. (**D**) Analysis of PSMG2-mNG2(11)-StrepII samples incubated at 37 °C for 1 h resulted in a cryo-EM map of a more mature 20S proteasome (pre-20S’). Scale bars: micrograph, 200 Å; 2D classes, 100 Å; cryo-EM map, 20 Å.

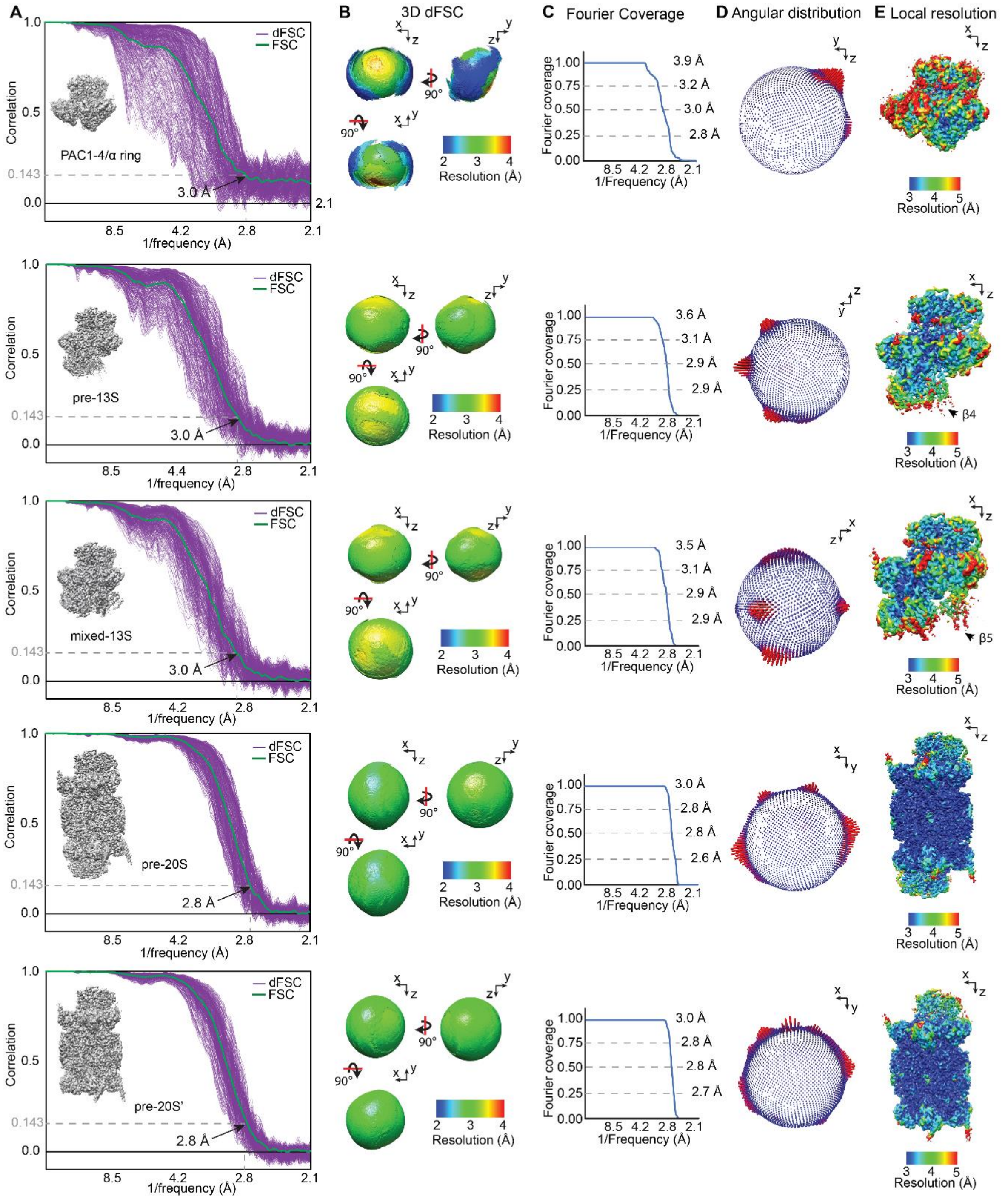

**Supplementary Figure 3. Cryo-EM map quality metrics**. (**A**) Fourier shell correlation (FSC) and 1D directional FSC (dFSC) plots. (**B**) Visualization of 3D dFSC. (**C**) Plots showing coverage of Fourier space. (**D**) Plots showing distribution of viewing angles from final refined datasets. (**E**) Local resolution maps.

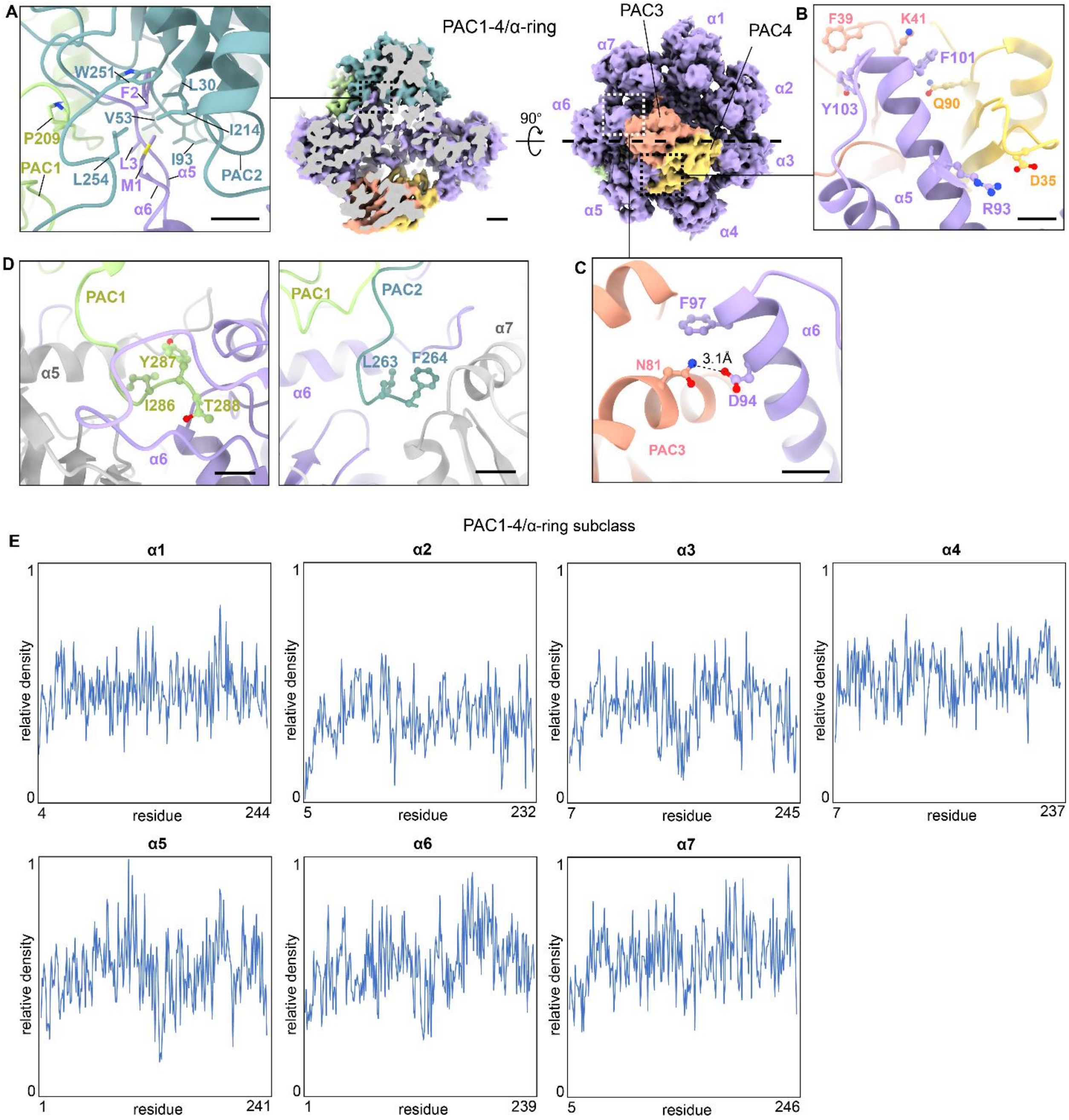

**Supplementary Figure 4. Interactions of α subunits with PAC1/PAC2 and PAC3/4.** (**A**) The N-terminal tails of α5 and α6 insert into a hydrophobic pocket between PAC1 and PAC2. Scale bar: 5 Å. (**B**) PAC3 and PAC4 clamp onto α5 through multiple sidechain interactions. Scale bars: atomic model, 5 Å; cryo-EM map, 10 Å. (**C**) PAC3 interaction with α6. Scale bar: 5 Å. (**D**) C-terminal tails of PAC1 and PAC2 in the HbYX-binding pockets. Scale bars: 5 Å. (**E**) Per residue cryo-EM density map values for each α subunit in the PAC1-4/α-ring subclass.

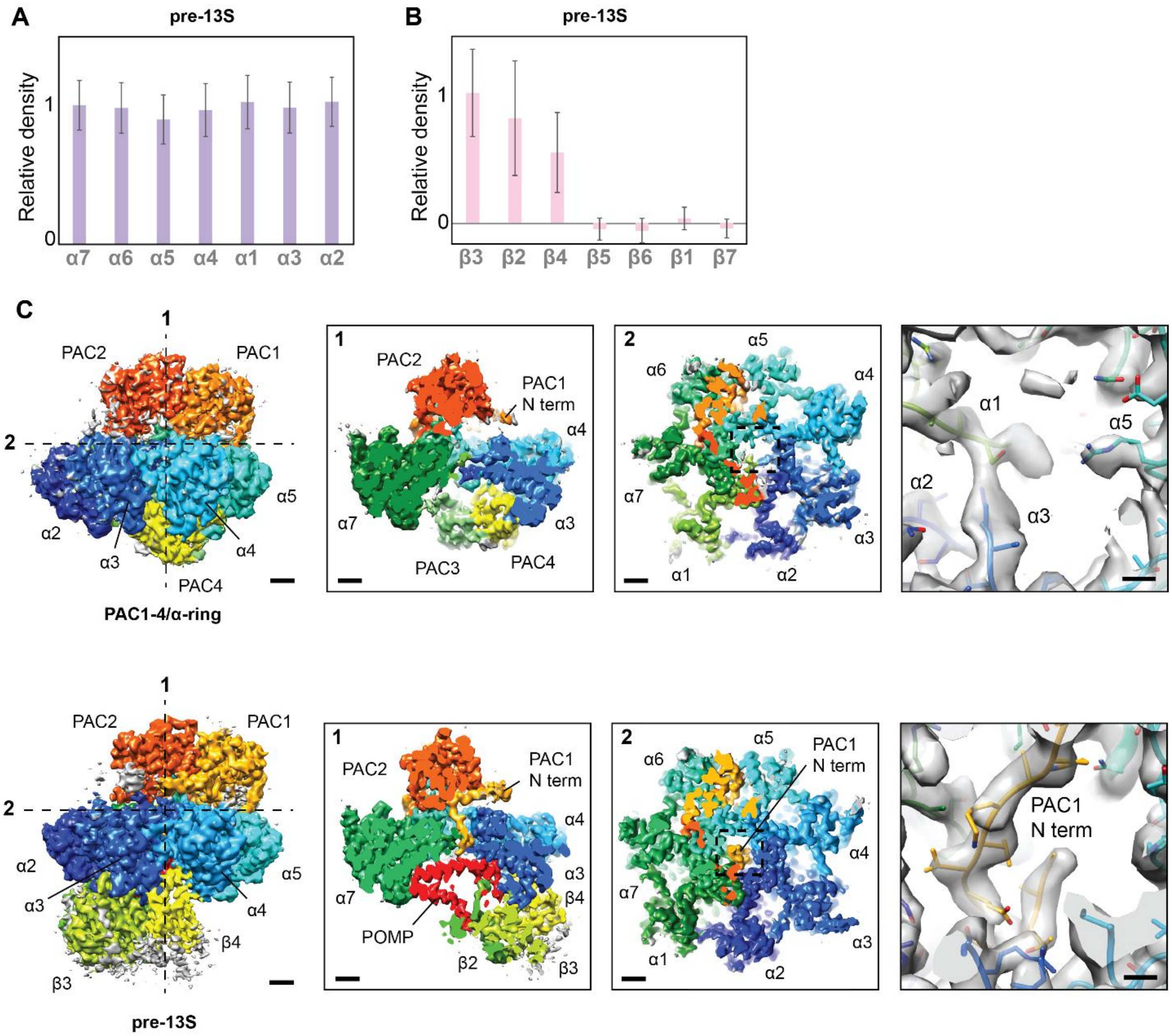

**Supplementary Figure 5. Analysis of the α-ring and β-ring of pre-13S.** (**A**) Quantification of the cryo-EM map density values of pre-13S reveal stoichiometric levels of each α subunit. (**B**) Quantification of the cryo-EM map density values of pre-13S β subunits reveals sub-stoichiometric levels of β4 and no binding of β1 and β5-7. (**C**) The cryo-EM map of pre-13S shows clear density for the N-terminal tail of PAC1 inserted into the α-ring pore, which is not observed in the PAC1-4/α-ring map. Scale bars: cryo-EM maps, 10 Å; atomic models, 2 Å.

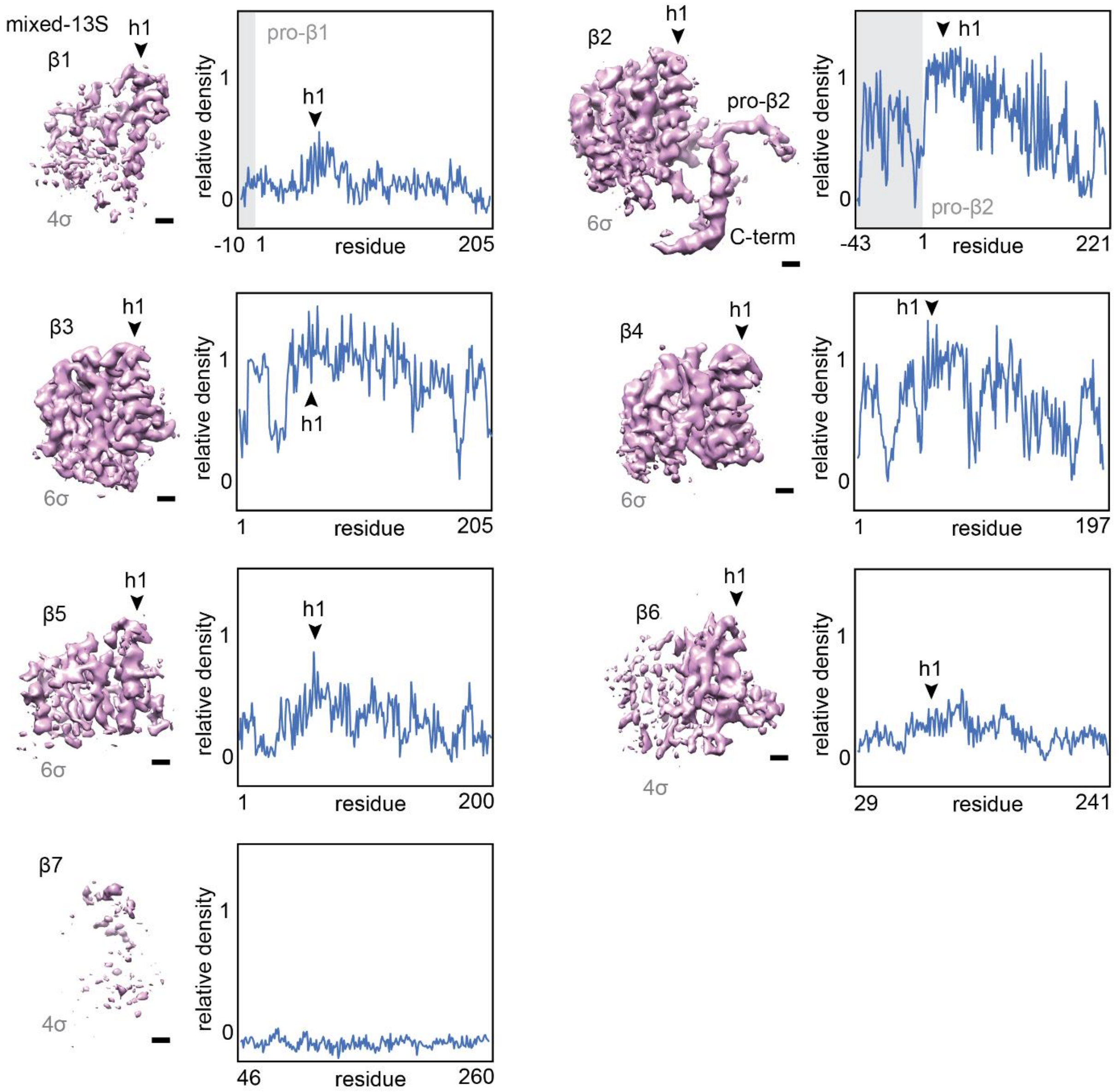

**Supplementary Figure 6. Cryo-EM density map analysis of the β subunits from the mixed-13S structure.** Density map of each subunit and corresponding per residue density values from the mixed-13S cryo-EM map. Strong density for helix 1 (h1) can be observed for the α-subunits. Scale bars: 5 Å.

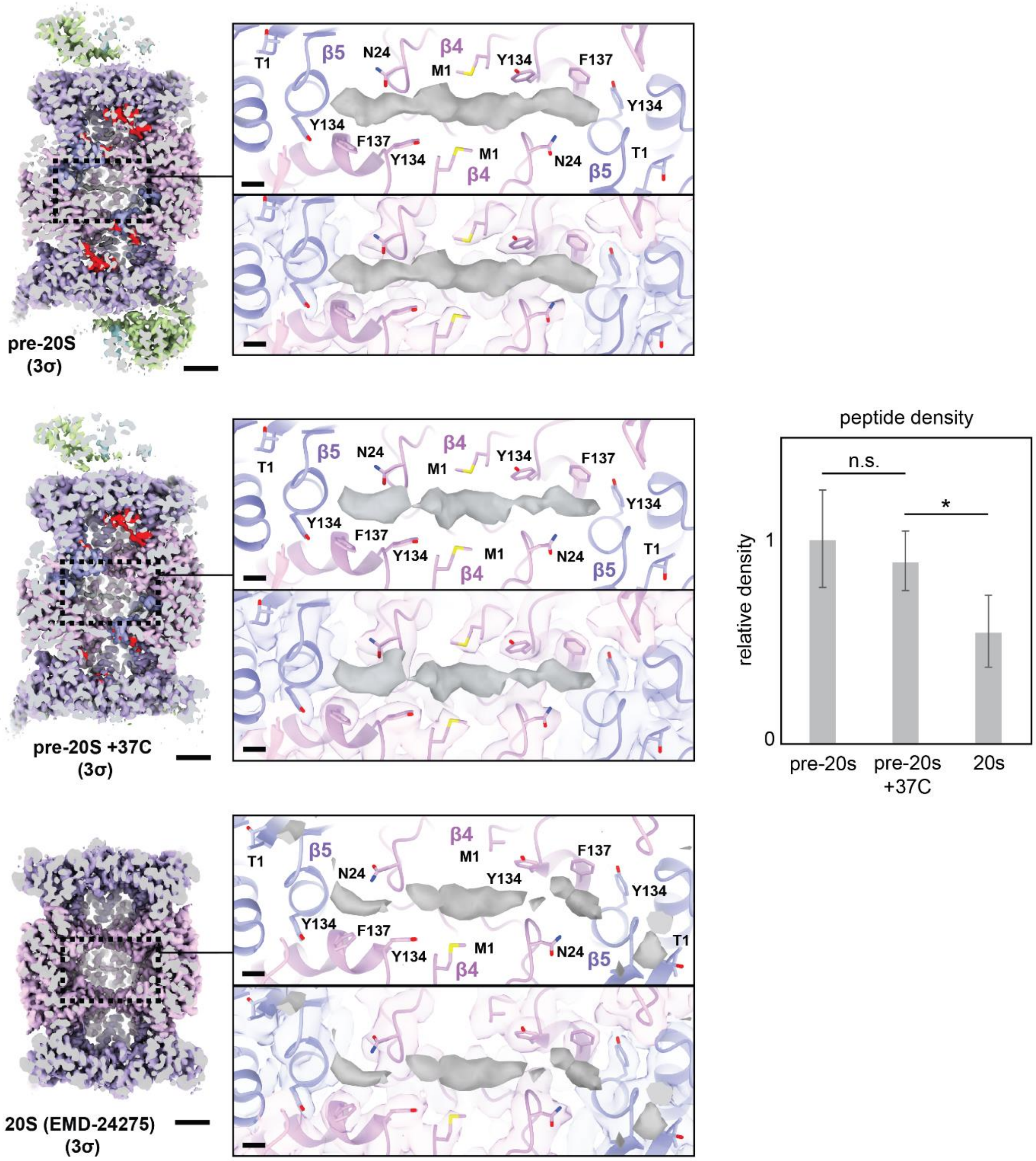

**Supplementary Figure 7. Unknown ligand bound at the dimer interface of two β4 subunits.** An extended strand of density is observed in the pre-20S cryo-EM map located in a groove between the β4 subunits, indicating the presence of an unknown ligand. The occupancy of the strand decreases relative to the β2 subunit as the 20S proteasome matures, but substantial levels remain in the mature 20S proteasome. The unknown ligand is close to β5 T1. Scale bars: cryo-EM map, 20 Å; atomic models, 3 Å. *p-value<0.001.

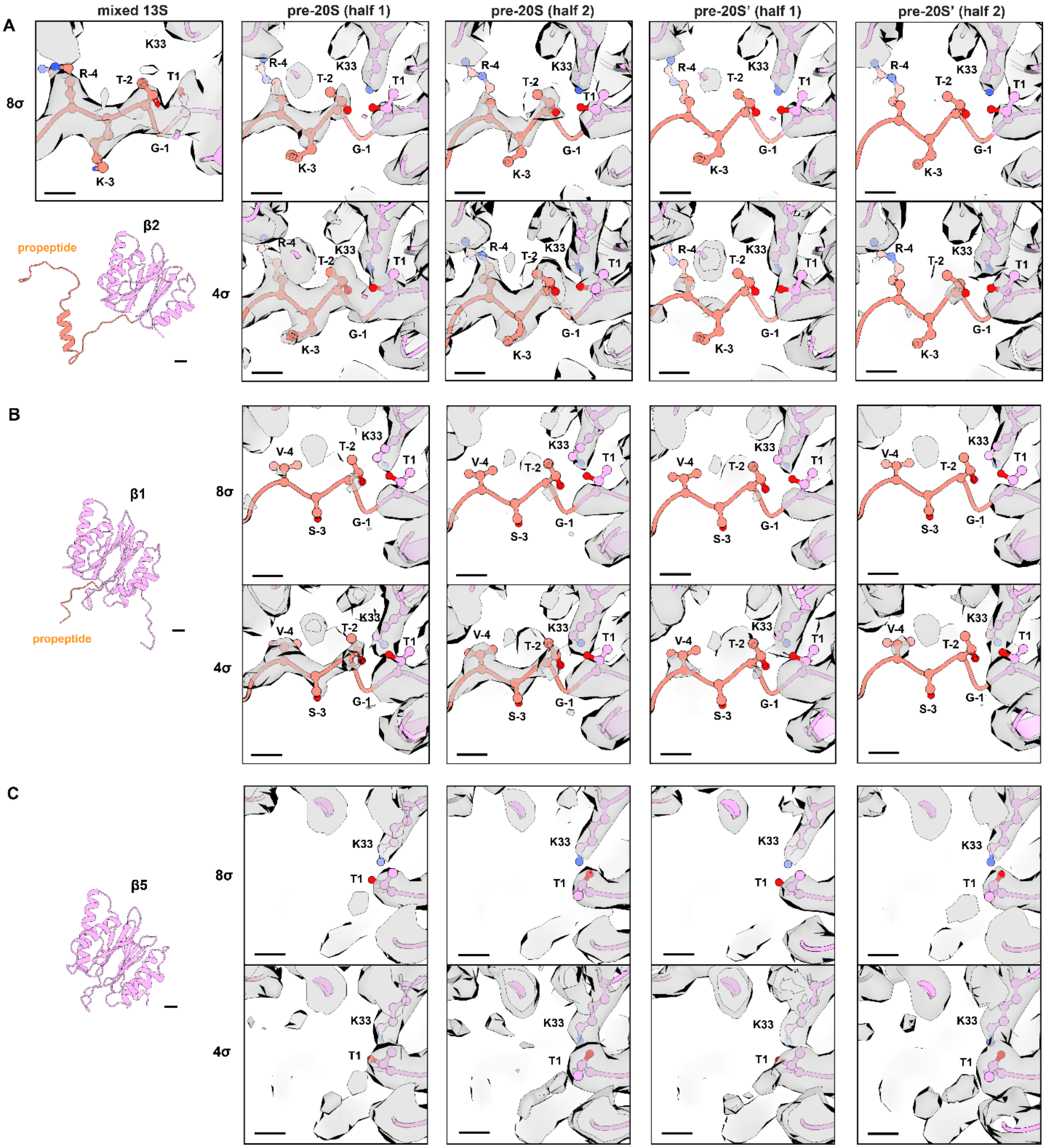

**Supplementary Figure 8. Cleavage of β pro-peptides.** (**A**) The β2 pro-peptide has higher density in the mixed-13S cryo-EM map and lower density in pre-20S and pre-20S' cryo-EM maps. Scale bars: subunit view, 5 Å; insets, 3 Å. (**B**) The β1 pro-peptide has low density in the pre-20S and pre-20S' cryo-EM maps, but density can still be observed at lower map thresholds (4σ). Scale bars: subunit view, 5 Å; insets, 3 Å. (**C**) No density is observable for the β5 pro-peptide. Scale bars: subunit view, 5 Å; insets, 3 Å.

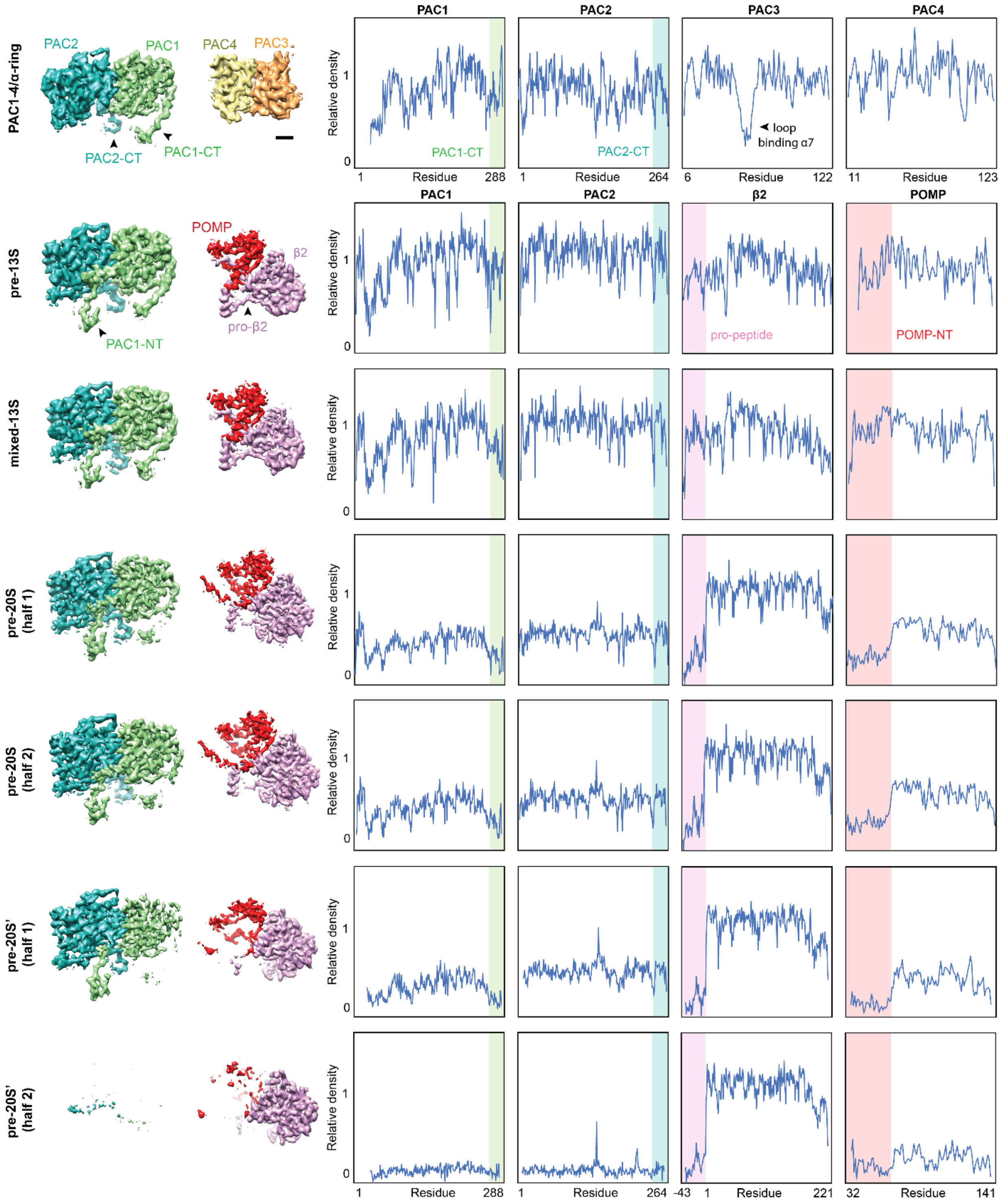

**Supplementary Figure 9. Cryo-EM map density analysis.** Isolated cryo-EM map densities for PAC1-4, POMP, and β2 across the different maps show sequential decreases in density for specific regions in the proteins as assembly progresses. Quantification of the map density on a per residue level shows clear boundaries where the density changes occur.

**Supplementary Table 1**. Cryo-EM data collection, processing, and validation parameters

| | PAC1-4/α-ring<br>EMDB-41377<br>PDB 8TM3 | Pre-13S<br>EMDB-41378<br>PDB 8TM4 | Mixed 13S<br>EMDB-41379<br>PDB 8TM5 | Pre-20S<br>EMDB-41380<br>PDB 8TM6 | Pre-20S'<br>EMDB-<br>41381 |
|---|---|---|---|---|---|
| **Data collection and processing** | | | | | |
| Magnification | 47,170 | 47,170 | 47,170 | 47,170 | 47,170 |
| Voltage (kV) | 300 | 300 | 300 | 300 | 300 |
| Electron exposure (e–/Å$^2$) | 30 | 30 | 30 | 30 | 30 |
| Defocus range (μm) | 1-2 | 1-2 | 1-2 | 1-2 | 1-2 |
| Pixel size (Å) | 1.06 | 1.06 | 1.06 | 1.06 | 1.06 |
| Symmetry imposed | C1 | C1 | C1 | C1 | C1 |
| Initial particle images (no.) | 3,782,868 | 3,999,264 | 3,999,264 | 882,852 | 1,877,850 |
| Final particle images (no.) | 111,057 | 57,026 | 44,644 | 115,008 | 100,657 |
| Map resolution (Å)<br>FSC threshold | 3.0<br>0.143 | 3.0<br>0.143 | 3.0<br>0.143 | 2.8<br>0.143 | 2.8<br>0.143 |
| Map resolution range (Å) | 2.8-3.9 | 2.9-3.6 | 2.9-3.5 | 2.6-3.0 | 2.7-3.0 |
| **Refinement** | | | | | |
| Initial model used (PDB code) | | | | | N/A |
| Model resolution (Å)<br>FSC threshold | 3.7<br>0.5 | 3.4<br>0.5 | 3.4<br>0.5 | 3.1<br>0.5 | N/A<br>N/A |
| Model resolution range (Å) | 2.9-3.7 | 2.9-3.4 | 2.9-3.4 | 2.7-3.1 | N/A |
| Map sharpening *B* factor (Å$^2$) | | | | | N/A |
| Model composition<br>Non-hydrogen atoms<br>Protein residues<br>Ligands | <br>17,563<br>2392<br>0 | <br>21,367<br>2806<br>0 | <br>21,689<br>2873<br>0 | <br>58,154<br>7687<br>0 | <br>N/A<br>N/A<br>N/A |
| *B* factors (Å$^2$)<br>Protein<br>Ligand | <br>116<br>N/A | <br>100<br>N/A | <br>100<br>N/A | <br>64<br>N/A | <br>N/A<br>N/A |
| R.m.s. deviations<br>Bond lengths (Å)<br>Bond angles (°) | <br>0.003<br>0.558 | <br>0.003<br>0.626 | <br>0.003<br>0.612 | <br>0.002<br>0.487 | <br>N/A<br>N/A |
| Validation<br>MolProbity score<br>Clashscore<br>Poor rotamers (%) | <br>1.72<br>7.48<br>0.17 | <br>1.63<br>7.37<br>0.22 | <br>1.58<br>7.11<br>0.09 | <br>1.27<br>4.20<br>0.12 | <br>N/A<br>N/A<br>N/A |
| Ramachandran plot<br>Favored (%)<br>Allowed (%)<br>Disallowed (%) | <br>95.53<br>4.38<br>0.09 | <br>96.50<br>3.46<br>0.04 | <br>96.90<br>3.03<br>0.07 | <br>97.71<br>2.23<br>0.05 | <br>N/A<br>N/A<br>N/A |
